# Machine learning-guided discovery of a conserved plasmid proteomic signature enables MALDI-TOF MS detection of pOXA-48-carrying *Enterobacterales*

**DOI:** 10.64898/2026.07.12.738056

**Authors:** Janko Sattler, Johannes Mueller-Reif, Dexiong Chen, Julian Sommer, Lucas Miranda, Juliette Murris, Till Hendrik Schulz, Yukino Gütlin, Janis Rogenmoser, Peter V. Treit, Thomas Pichl, Ela Sauerborn, Helena M. B. Seth-Smith, Tim Roloff, Stephan Göttig, Jonathan Jantsch, Andreas F. Wendel, Jacob Moran-Gilad, Matthias Mann, Axel Hamprecht, Adrian Egli, Karsten Borgwardt

## Abstract

OXA-48 carbapenemases are among the most widespread and important resistance mechanisms in *Enterobacterales*. Yet detecting carbapenemases by conventional workflows necessitates additional testing, thus delaying optimization of therapy and implementation of infection control measures. Here, we present a machine learning approach that identifies the conserved pOXA-48 plasmid directly from routine MALDI-TOF spectra acquired for species identification. The model detects pOXA-48 carriers with an AUROC of 0.96–0.98 across two independent hospital cohorts and instrument platforms, indicating near-perfect discrimination. Using bottom-up proteomics, plasmid conjugation, and plasmid curing, we link the discriminative MALDI-TOF spectral features to proteins encoded on pOXA-48, with DUF1496 domain-containing protein producing the most discriminative spectral feature. Our approach reframes the resistance prediction task from inferring a resistance phenotype to detecting a conserved plasmid through its expressed proteomic signature and has the potential to enable rapid MALDI-TOF MS-based diagnostics for a wide range of plasmid-based resistance determinants.

## Introduction

Antimicrobial resistance is one of the biggest threats to global healthcare, directly attributable to over a million deaths annually ^1,2^. Among Gram-negative bacteria, carbapenemase-producing *Enterobacterales* represent a major challenge for treatment and infection prevention and control as they are highly prevalent ^3^, resistant to many last-line antibiotics, and spread efficiently via horizontal gene transfer ^4^. Carbapenemases are enzymes that can hydrolyse carbapenems and most other β-lactam antibiotics, rendering their use ineffective. Among these enzymes, OXA-48-like carbapenemases are globally among the three most prevalent carbapenemase types in *Enterobacterales* ^5,6^, and have emerged as the dominant carbapenemase type across parts of Europe, North Africa, and the Middle East, where they are now endemic ^7,8^. This is of high clinical importance, as bloodstream infections with OXA-48-producing *Enterobacterales* carry a 30-day mortality of up to 50% ^9^. Furthermore, unlike most other carbapenemases, OXA-48-like carbapenemases are characterized by limited hydrolytic activity against extended-spectrum cephalosporins, frequently resulting in subtle or ambiguous resistance phenotypes that complicate detection by routine susceptibility testing ^10^. Consequently, OXA-48-like-producing isolates often evade early recognition, facilitating silent dissemination.

The spread of OXA-48, the most common OXA-48-like variant, and the single amino acid derivative OXA-162, is driven predominantly by a highly conserved IncL plasmid backbone, pOXA-48, which carries *bla*_OXA-48_ or *bla*_OXA-162_ in a remarkably stable genetic context ^7,11,12^. This plasmid exhibits high conjugation efficiency and an unusual capacity to cross species barriers within *Enterobacterales* ^11,13,14^. As a result, genetically diverse hosts can acquire near-identical plasmids, decoupling resistance spread from clonal expansion, and further contributing to global dissemination.

Matrix-assisted laser desorption/ionization time-of-flight mass spectrometry (MALDI-TOF MS) is the standard platform for bacterial species identification in clinical microbiology, routinely applied to colonies recovered by primary culture. While species-level results are available at this earliest diagnostic time point, resistance type classification typically requires further testing such as phenotypic antimicrobial susceptibility testing (AST) and confirmatory testing for carbapenemase detection. For OXA-48/-162-producers, whose subtle phenotypes frequently fail to trigger suspicion even after AST results, this delay creates a critical window during which carriers go unrecognized, undermining infection prevention and control as well as targeted therapy ^9^.

MALDI-TOF spectra contain protein signals beyond those used for taxonomy. However, machine-learning approaches to extract resistance information have so far relied on correlations between MALDI-TOF spectra and AST data from discrete isolate collections, lacking interpretability and robustness outside the training setting, in part because the same phenotype can arise from genetically distinct mechanisms. Direct detection of the OXA-48/-162 enzyme is not feasible at routine MALDI-TOF settings, as the singly-charged enzyme exceeds the default 2–20 kDa analytical range ^15^. However, prior work on other carbapenemases suggests plasmid-encoded accessory proteins can act as surrogate markers ^16^, and the exceptional conservation of pOXA-48 makes it a promising candidate for such an approach. We therefore reframe the prediction task from inferring a resistance phenotype to detecting a conserved plasmid through its expressed proteomic signature. We hypothesized that pOXA-48-encoded proteins co-transferred with *bla*_OXA-48/-162_ generate reproducible spectral signatures exploitable for a robust and interpretable detection.

Here, we present a biology-driven framework for detecting pOXA-48-carrying *Enterobacterales* from standard MALDI-TOF MS spectra, flagging probable *bla*_OXA-48/OXA-162_ carriers at the point of species identification rather than after conventional resistance testing (Fig. 1a). By integrating microbial genomics, experimental plasmid biology, and machine learning, we move beyond black-box prediction and systematically link discriminative spectral features to defined plasmid-encoded proteins (Fig. 1b). Using plasmid conjugation and curing experiments, bottom-up proteomics, and examination of naturally occurring pOXA-48 truncation variants, we establish causal relationships between specific MALDI-TOF peaks and the presence of the pOXA-48 plasmid (Fig. 1c). We further demonstrate that these biologically grounded features enable robust prediction across species, acquisition protocols, and instrumentation. Together, our results illustrate how mechanistic insight into microbial genetics can be leveraged to transform MALDI-TOF-based resistance prediction from empirical pattern recognition into an interpretable strategy whose expected generalizability is grounded in conserved plasmid biology.

**Fig. 1.**
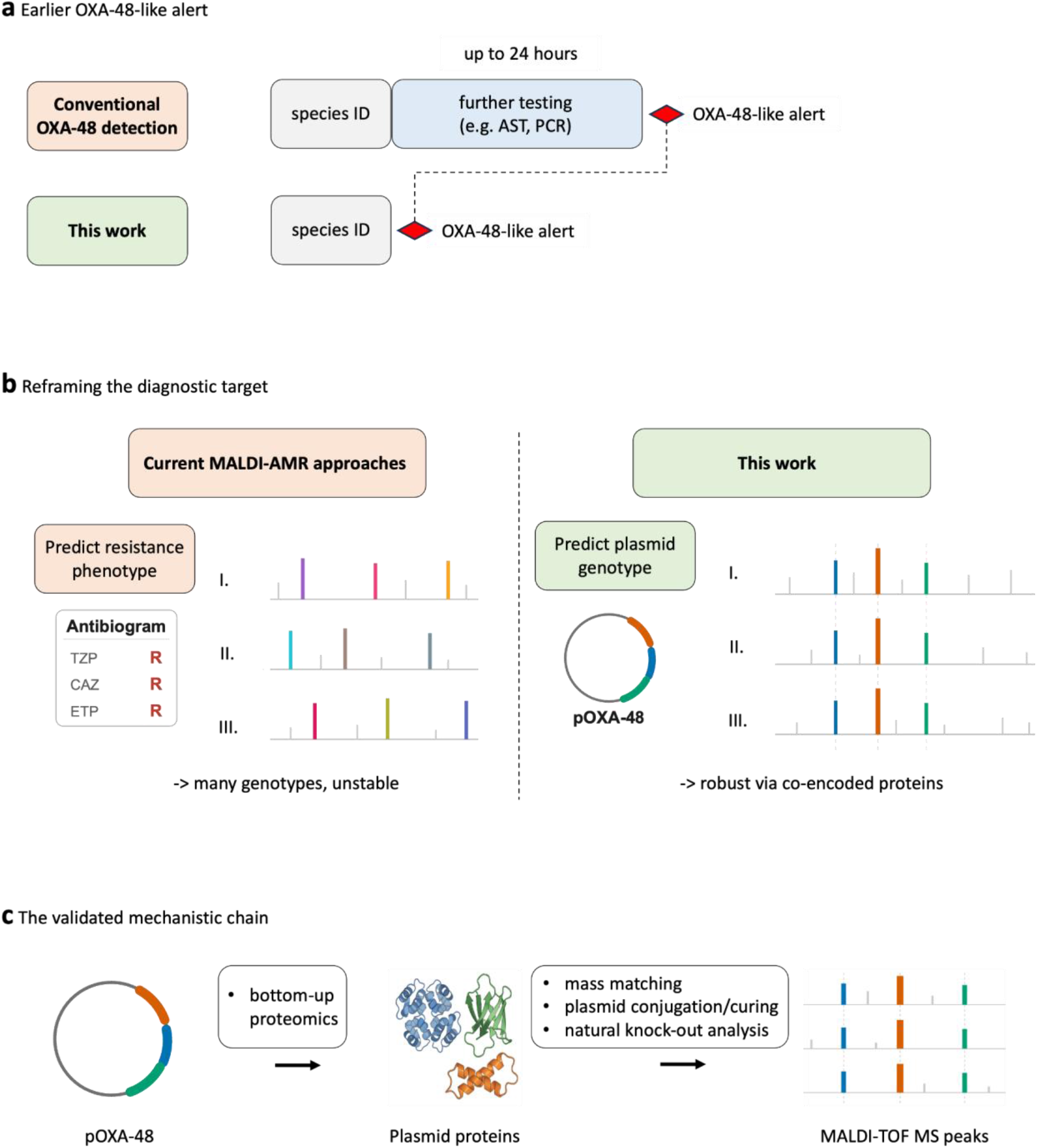
A plasmid-targeted strategy for earlier, more robust MALDI-TOF detection of OXA-48/-162-producing *Enterobacterales*. **a**, Time-to-result in the diagnostic workflow. In the conventional workflow (top), MALDI-TOF species identification is followed by additional testing such as overnight phenotypic antimicrobial susceptibility testing (AST) and/or confirmatory OXA-48-like testing before an OXA-48-like alert (red diamond) is raised. In the approach proposed here (bottom), the alert is raised already during the species identification up to 24 h earlier. **b**, Two contrasting machine-learning targets. Left (“Current MALDI-AMR approach”): a model trained to predict a resistance phenotype. The highlighted discriminative peaks (coloured) are drawn at different m/z in each of three example spectra. Right (“This work”): the conserved pOXA-48 (IncL) plasmid (ring, with colour-coded genes) as target. Three coloured peaks corresponding to co-encoded plasmid proteins align at the same m/z across three example spectra (dashed lines), despite the variable chromosomal background (grey). Each stacked trace represents a spectrum from a different isolate or species. **c**, The mechanistic chain validated in this study: the pOXA-48 plasmid encodes proteins whose expression produces discriminative MALDI-TOF MS peaks. Labels beneath the arrows indicate the experimental and computational methods used to establish each link: bottom-up proteomics connects the plasmid to its expressed proteins, and mass matching, plasmid conjugation and curing, and analysis of a naturally occurring pOXA-48 deletion variant connect those proteins to the spectral peaks.

## Results

### Commercial software for MALDI-TOF diagnostics fails to identify OXA-48/-162-producing *Enterobacterales*

To establish a diagnostic baseline, we evaluated Clover MS Data Analysis Software, to our knowledge the only commercially available system for detecting OXA-48/-162 carbapenemases from standard MALDI-TOF spectra, based on the approach of Gato et al. ^17^. We applied it to 166 *Klebsiella pneumoniae* complex isolates from our internal datasets. The binary carbapenemase detection model achieved perfect specificity (100%) but near-complete failure in sensitivity: only 1 of 83 OXA-48/-162-positive isolates was identified as carbapenemase-positive (sensitivity 1.2%, Extended Data Table 1), based on 2 of 3 technical replicates yielding a positive prediction. When this isolate was further classified by the subtype module, one replicate was assigned to OXA-48 and one to KPC, indicating unreliable subtype discrimination even among correctly detected cases. These results show that robust OXA-48/-162 detection from standard MALDI-TOF spectra remains challenging, motivating us to develop a machine-learning approach informed by plasmid biology.

### A mixed-protocol machine learning framework achieves robust cross-domain and cross-instrument detection

To determine whether MALDI-TOF spectra contain sufficient information for robust pOXA-48 detection, we developed a supervised classification framework trained on spectra acquired under multiple conditions. The dataset comprised 246 ‘in-domain’ isolates measured under a standard acquisition protocol and 60 ‘cross-domain’ isolates measured under four additional conditions designed to test robustness: i) omission of ertapenem pre-selection; ii) omission of formic acid overlay; iii) culture on Mueller-Hinton agar; and iv) measurement from an alternative blood agar (Methods; Fig. 2a).

**Fig. 2.**
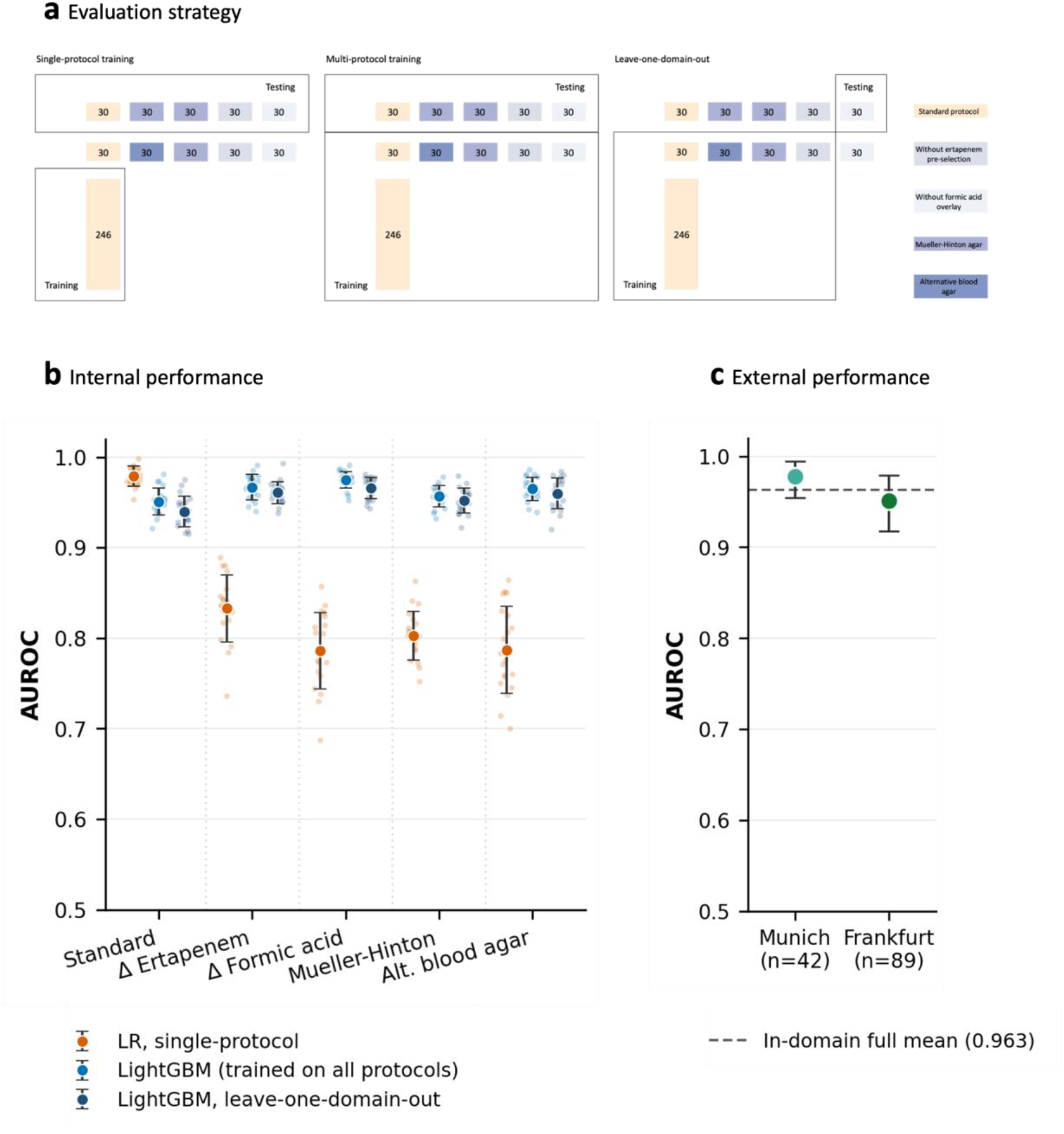
A multi-protocol machine-learning framework enables robust cross-domain and cross-instrument detection of pOXA-48. **a**, Evaluation strategy. Three set-ups: single-protocol training on standard-protocol spectra; multi-protocol training incorporating cross-domain isolates with inverse-proportional condition weighting; and leave-one-domain-out (LODO) validation, in which each acquisition protocol is sequentially excluded from training and used only for evaluation. Numbers indicate isolate counts per block; rows correspond to the five acquisition protocols (legend, right). **b**, Internal cross-domain classification performance. AUROC for pOXA-48 detection across the five acquisition protocols for three models: LASSO logistic regression trained on standard-protocol spectra only (vermillion), the multi-protocol LightGBM model trained on temporally augmented, condition-weighted data from all five protocols (blue), and the LODO LightGBM model (dark blue). Points, mean AUROC across 20 random seeds, with the 20 individual per-seed values overlaid as dots; error bars, ±1 s.d. **c**, External classification performance. AUROC of the LightGBM full model (6,004 features) on two independent cohorts: Munich (n = 42; Bruker Biotyper) and Frankfurt (n = 89; bioMérieux VITEK MS). Points are point estimates (mean of 100 Monte Carlo spectrum draws on the full cohort). Error bars are 95% confidence intervals from an isolate-level Monte Carlo-aware percentile bootstrap (10,000 resamples, stratified by class). Dashed line, mean in-domain AUROC of the full model across the five acquisition protocols (0.963). Numerical values are given in Extended Data Tables 2 and 3.

A LASSO logistic regression model trained exclusively on spectra acquired under the standard protocol achieved strong in-domain performance in distinguishing spectra of isolates positive for pOXA-48 from negative ones (area under the receiver operating characteristic curve, AUROC = 0.979 ± 0.011). However, performance dropped when the model was applied to spectra generated under altered acquisition conditions, with AUROC decreasing to 0.786–0.833 across cross-domain protocols (Fig. 2b; Extended Data Table 2). This indicates that models trained on a single experimental protocol are highly sensitive to the experimental variability commonly encountered in routine clinical workflows.

To improve cross-domain performance, we applied temporal augmentation, inverse-proportional condition weighting (Methods), and used LightGBM as classification algorithm, all of which independently improved performance and exhibited additive effects (Extended Data Table 2). The resulting model, trained on the combined multi-protocol and temporally augmented dataset (LightGBM-full), achieved AUROC values of 0.951–0.975 across all five acquisition protocols, largely recovering the performance loss observed for single-protocol models (Fig. 2b). Performance was stable across a set of LightGBM hyperparameter configurations (mean AUROC 0.958–0.968; Supplementary Table 1). Consistent results were obtained in a leave-one-domain-out analysis, in which each acquisition protocol was sequentially excluded from training and used exclusively for evaluation (AUROC 0.940–0.966; Extended Data Table 2), indicating that the generalizability gain of the augmentations can also remain in unseen conditions.

To further test generalization across instrumentation, we evaluated the LightGBM-full model on two independent spectra collections of clinical isolates from the Technical University of Munich (n = 42) and from University Hospital Frankfurt (n = 89). These isolates were cultured using the same protocol and were measured either on a Bruker Biotyper (Munich) or on a bioMérieux VITEK MS system (Frankfurt). Without retuning, the model achieved an AUROC of 0.978 (95% CI 0.954–0.995) on the Munich dataset and of 0.951 (95% CI 0.918–0.979) on the Frankfurt dataset, indicating cross-instrument transferability (Fig. 2c, Extended Data Table 3).

### A multi-peak plasmid signature underlies classification performance

To identify the spectral features driving classification, we computed permutation feature importance for the LightGBM-full model across all 20 seeds on the internal test sets (Methods). Importance was distributed across multiple m/z regions rather than concentrated in a single peak (Fig. 3a). Although the m/z 10,163 bin ranked highest, additional positions contributed substantially.

**Fig. 3.**
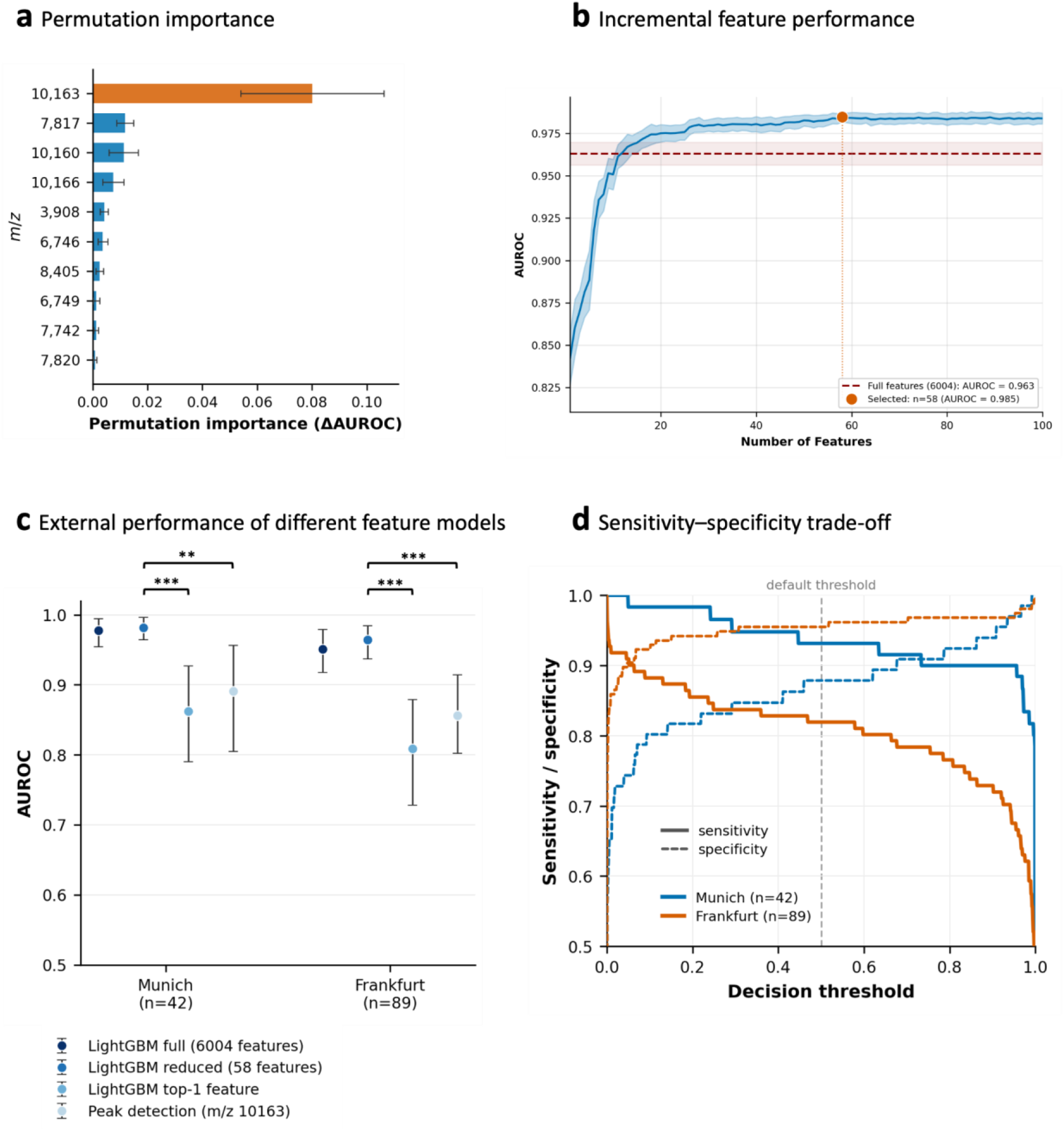
A compact multi-peak feature set drives classification and outperforms single-feature detectors. **a**, Permutation feature importance of the ten highest-ranked features, computed on internal cross-domain validation data as the mean decrease in AUROC over 20 random seeds. The feature at m/z 10,163 is shown in vermillion and the remaining nine in blue; error bars, ±1 s.d. across seeds; annotated values are feature m/z positions. **b**, AUROC as a function of the number of features added in order of decreasing permutation importance. Blue line, mean AUROC across 20 seeds (shaded band, ±1 s.d.); dark-red dashed line, AUROC of the full-feature (6,004) model (band, ±1 s.d.); vermillion circle, the feature count selected by the one-standard-error rule (n = 58). **c**, External AUROC on the Munich (n = 42) and Frankfurt (n = 89) cohorts for four classifiers of decreasing complexity (sequential blue gradient, darker = more complex): LightGBM full (6,004 features), LightGBM reduced (58 features), LightGBM restricted to the single top-ranked feature (the m/z 10,163 bin), and a rule-based peak detector at m/z 10,163. Error bars are 95% confidence intervals (isolate-level Monte Carlo-aware bootstrap, 10,000 resamples, stratified by class). Differences between the LightGBM-reduced model and the LightGBM top-1 feature model and between the LightGBM-reduced model and the single peak detection model are statistically significant; asterisks indicate P value (* ≤ 0.05, **≤ 0.01, ***≤ 0.001) (Methods). Numerical values are given in Extended Data Table 3. **d**, Sensitivity (solid) and specificity (dashed) of the LightGBM-reduced model as a function of the decision threshold, for the Munich (n = 42; blue) and Frankfurt (n = 89; vermillion) cohorts; each curve is the mean of 100 Monte Carlo spectrum draws (one replicate per isolate). The dashed vertical line marks the default 0.5 threshold (values in Extended Data Table 3).

Guided by this ranking, we performed sequential feature reduction to determine the minimal feature set required for robust classification (Methods). The highest mean performance (AUROC = 0.985) was achieved with a model using 58 features (Fig. 3b, Extended Data Fig. 1), hereafter referred to as LightGBM-reduced. LightGBM-reduced matched or slightly exceeded the full model on the independent external Munich (AUROC = 0.982 versus 0.978) and Frankfurt datasets (AUROC = 0.964 versus 0.951), despite feature selection being performed exclusively on internal data (Fig. 3c, Extended Data Table 3). This indicates that importance-guided feature reduction removes instrument- or site-specific noise while retaining biologically generalizable signal. Per-species AUROC remained ≥ 0.93 in every external stratum, so aggregate performance is not driven by cohort species composition (Extended Data Tables 4, 6). Differences in sensitivity and specificity at the 0.5 decision threshold reflect calibration shifts between feature sets, while threshold-independent ranking quality is preserved. These operating characteristics could be adjusted for a screening use case: lowering the decision threshold below the 0.5 default reached a ∼95% sensitivity operating point at a specificity of 0.83 (Munich) and 0.78 (Frankfurt) (Fig. 3d).

To test whether the dominant feature alone could explain classification performance, we pursued two complementary single-feature approaches on both external datasets. A LightGBM model restricted to the single most important binned feature (LightGBM-top1) achieved AUROC values of 0.862 and 0.809 for Munich and Frankfurt, respectively. A rule-based single-peak detection classifier operating on unbinned preprocessed spectra and assigning OXA-48/-162-positive status based on detection of a peak at m/z 10,163 (with optimized parameters; Methods), achieved AUROC values of 0.891 and 0.856. Both approaches performed significantly worse than the LightGBM-reduced model (AUROC = 0.982 and 0.964 for Munich and Frankfurt; Extended Data Table 3). These results confirm that robust pOXA-48 detection requires the full multi-peak signature, consistent with a model that exploits multiple co-encoded plasmid proteins rather than a single biomarker.

### The LightGBM-reduced model generalizes to clinically relevant edge cases

The LightGBM-reduced model was tested internally on two clinically relevant subsets that were excluded from training: isolates co-producing a second carbapenemase with species-matched negative controls (predominantly NDM co-producers) and OXA-48/-162-carrying species not represented in training with species-matched negative controls. For double-carbapenemase producers, the model achieved an AUROC of 0.901, indicating that co-harboured resistance determinants do not substantially interfere with pOXA-48 detection. Across rare species, the model achieved an aggregated AUROC of 0.832. Performance varied by species, reaching values comparable to the training species for *Morganella morganii* (AUROC 0.990) and *Serratia marcescens* complex (AUROC 0.890) (Extended Data Table 5).

### Horizontal gene transfer and plasmid curing establish pOXA-48 as the signal source

To test the biological origin of the classifier’s multi-peak signature, we performed isogenic gain- and loss-of-function experiments. Transfer of pOXA-48 from clinical donor MMC-K-0255 into the plasmid-free recipients *Escherichia coli* J53 and *Klebsiella quasipneumoniae* PRZ ^18^ via conjugation reverted both recipients from OXA-48-negative classifications (probability = 0.14 and 0.00, respectively) to OXA-48-positive (both 1.00), with concomitant emergence of the dominant peak at m/z 10,163 (Fig. 4a, 4b). Reciprocal removal of pOXA-48 from clinical isolate MMC-K-0225 by plasmid curing reverted the classification from positive (0.78) to negative (0.00), and the m/z 10,163 peak disappeared (Fig. 4c). This bidirectional gain and loss across two species supports that classification tracks pOXA-48 carriage itself, not chromosomal background, clonal lineage, or technical artefacts.

**Fig. 4.**
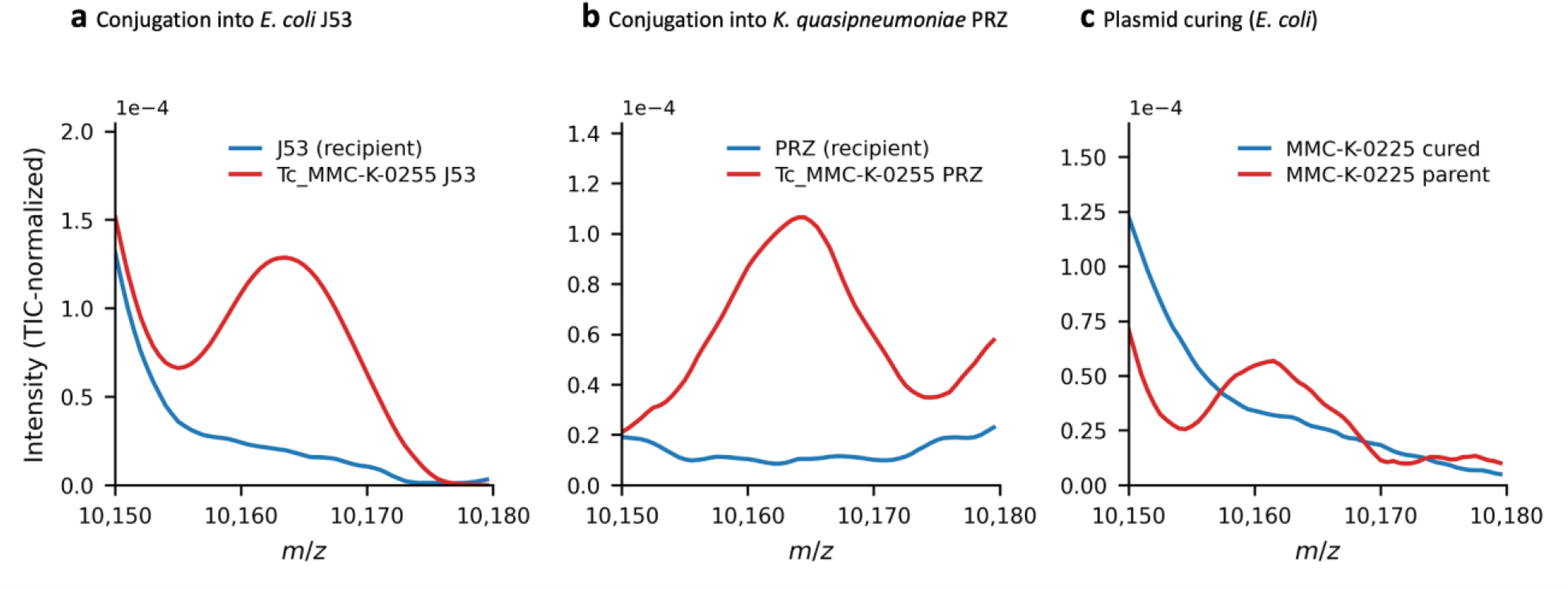
Conjugation and plasmid curing identify pOXA-48 as the source of the spectral signature. MALDI-TOF spectra (mean of three technical replicates, total-ion-current–normalized) in the m/z 10,150–10,180 region. **a**, Plasmid-free *Escherichia coli* J53 (blue) and the transconjugant carrying pOXA-48 from clinical donor MMC-K-0255 (red). **b**, Plasmid-free *Klebsiella quasipneumoniae* PRZ (blue) and the transconjugant carrying the same plasmid (red). **c**, Clinical isolate *E. coli* MMC-K-0225 carrying pOXA-48 (red, parent) and after plasmid curing (blue, cured). Throughout, blue indicates the absence and red the presence of pOXA-48.

### *In silico* mass prediction and bottom-up proteomics map discriminative spectral features to pOXA-48-encoded proteins

To map the classifier’s discriminative features to defined molecular species, we performed label-free bottom-up proteomics on 17 clinical isolates (8 pOXA-48-positive, 9 negative) spanning three *Enterobacterales* species (*K. pneumoniae*, *E. coli*, *Enterobacter cloacae* complex), sequence-type-matched where feasible (Supplementary Table 3). Of the 81 intact pOXA-48 coding sequences (CDS), 58 yielded peptide-level evidence and were detected almost exclusively in pOXA-48-positive isolates (Fig. 5a, 5b; Supplementary Tables 4 and 5). Low-level signal observed in pOXA-48-negative isolates (Fig. 5a) is characteristic of library-driven peptide propagation in data-independent acquisition (DIA) proteomics, with ∼24-fold fewer precursors detected per negative isolate than per positive (Supplementary Table 4). Chromosomally-encoded proteins, in contrast, showed no class discrimination (Extended Data Fig. 2), confirming that the differential signal arises specifically from plasmid-borne gene products.

**Fig. 5.**
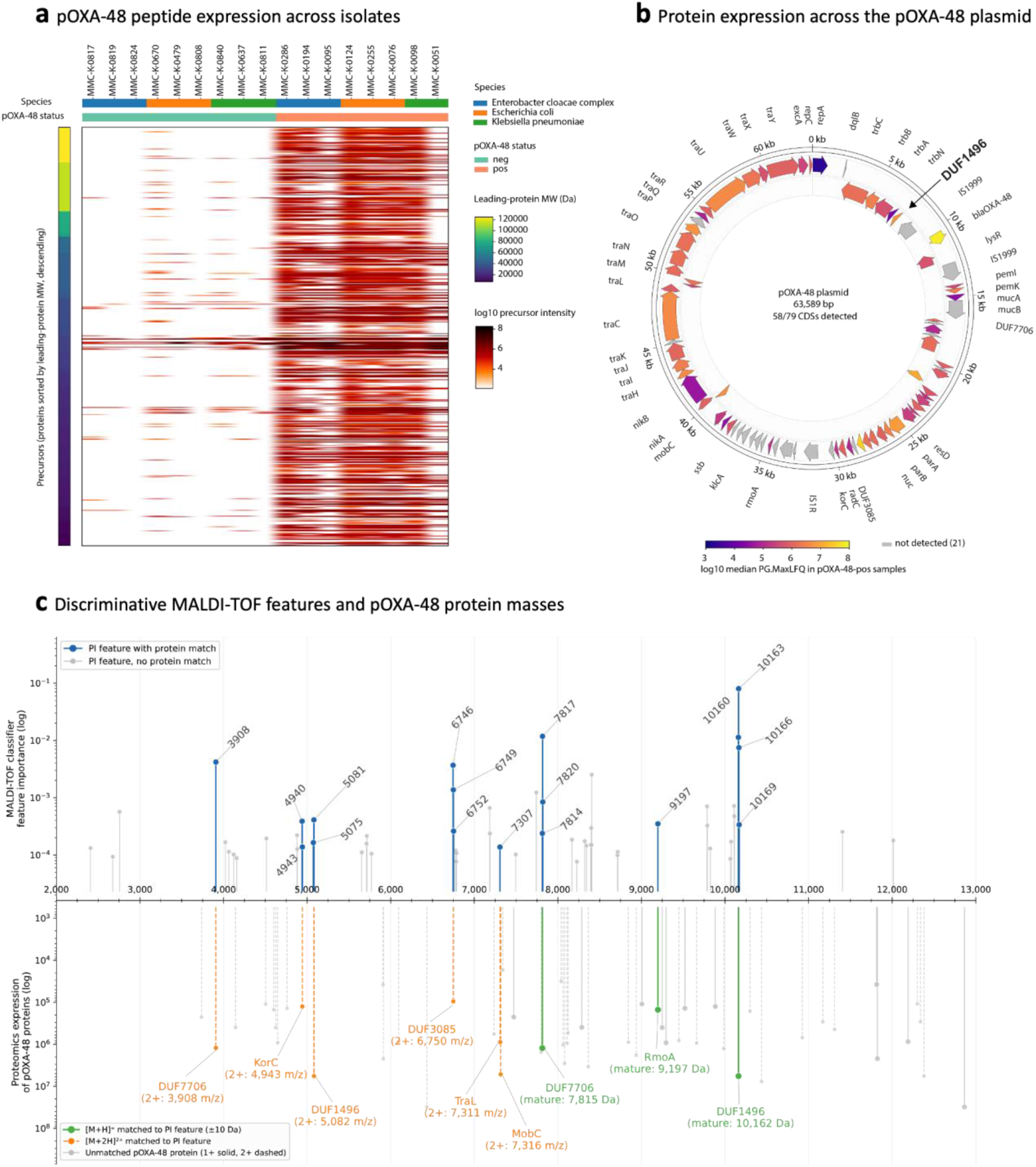
Discriminative MALDI-TOF features correspond to expressed pOXA-48-encoded proteins. **a**, Heatmap of pOXA-48-derived peptide precursor intensities (log10 normalized intensity; 0.1% FDR). Rows, 597 precursors from 58 pOXA-48-specific protein groups, grouped by protein and ordered by descending full-length molecular weight of the leading protein (left colour bar, viridis; ∼7.5–125 kDa); white lines separate protein groups. Columns, 17 isolates (9 pOXA-48-negative, 8 pOXA-48-positive; top colour bars indicate species and pOXA-48 status. **b**, Circular map of the pOXA-48 plasmid (63,589 bp). The 79 rendered CDS are shown on the outer (forward strand) and inner (reverse strand) tracks (Methods); labels mark annotated genes. Of these 79 CDS, 58 were detected and are coloured by log10 median label-free quantification intensity across the 8 pOXA-48-positive isolates (plasma colourmap); the remaining 21 are shown in grey. **c**, Correspondence between discriminative MALDI-TOF features and pOXA-48 protein masses. Top, permutation importance of the 58 retained features placed on the m/z axis; features with a pOXA-48 protein mass within ±10 Da (singly- or doubly-charged ion) are blue, unmatched features grey, and matched features are labelled by m/z bin index. Bottom, predicted average masses of the 58 detected pOXA-48 proteins, placed at the m/z of their singly-charged (solid stems) or doubly-charged (dashed stems) ion; green, matched to a feature as [M+H]⁺; orange, matched as [M+2H]²⁺; grey, unmatched. Mature masses (after predicted signal-peptide cleavage) are shown where a signal peptide was predicted, full-length masses otherwise.

Cross-referencing the predicted average masses of the 58 detected pOXA-48 proteins against the 58 retained ML features yielded protein-level matches for 17 features (Fig. 5c). This is a high proportion, as most detected proteins fall outside the standard MALDI-TOF mass range, and predicted masses do not account for post-translational modifications that can shift a protein away from its expected m/z. The 17 matches exceeded every one of 10,000 random feature placements (median 5, maximum 15; empirical P < 0.0001), supporting that the classifier’s discriminative signal corresponds to defined, expressed pOXA-48 gene products rather than incidental spectral correlations. Strikingly, eight of the ten most informative features mapped to only three proteins: a DUF1496 domain-containing protein (ranks 1, 3, 4; singly-charged ions of the mature 10.2-kDa form), a DUF7706 domain-containing protein (ranks 2, 5, 10; singly-and doubly-charged ions of the 7.8-kDa protein), and a DUF3085 domain-containing protein (ranks 6, 8; doubly-charged ions of the 13.5-kDa protein). The DUF1496 and DUF7706 proteins were also among the most abundant pOXA-48-encoded proteins in the MALDI-TOF mass range (Fig. 5b, 5c), consistent with a signature dominated by a small number of highly expressed, plasmid-conserved gene products.

### The m/z 10,163 peak depends on the intact DUF1496 protein

To directly link the dominant feature to the DUF1496 protein, we electroporated *E. coli* J53 with a natural pOXA-48 variant carrying a pOXA-48 truncation around the gene encoding the DUF1496 protein (donor MMC-K-0151, Extended Data Fig. 3)^19^. Whereas conjugation of the full pOXA-48 from MMC-K-0255 produced a prominent m/z 10,163 peak (Fig. 6a, red), the transformant carrying the truncated plasmid showed no discernible peak (Fig. 6a, orange) and was indistinguishable from the plasmid-free recipient at this m/z position (Fig. 6a, blue). This single-gene-resolved comparison establishes the DUF1496 protein as the molecular correlate of the dominant spectral feature.

**Fig. 6.**
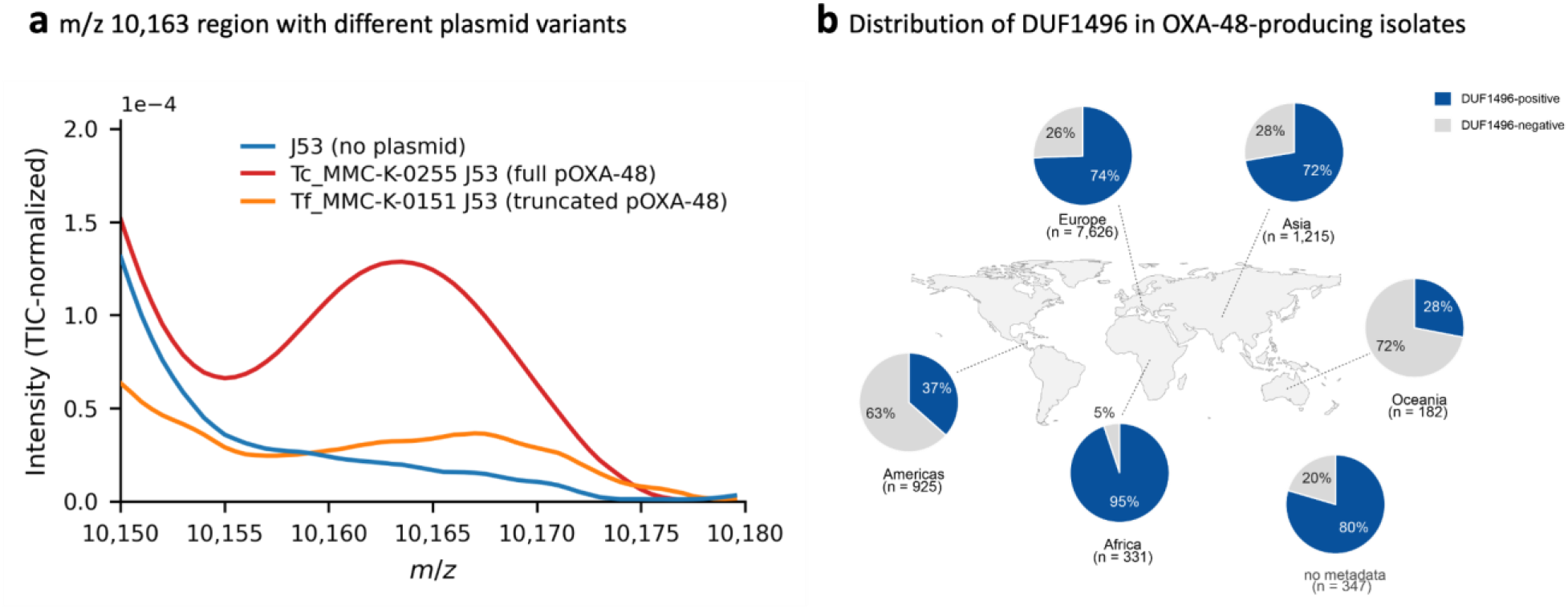
The dominant m/z 10,163 peak depends on the DUF1496 protein, which is highly conserved across pOXA-48 worldwide. **a**, MALDI-TOF spectra (mean of three technical replicates, total-ion-current–normalized; m/z 10,150–10,180) of three *Escherichia coli* J53-derived strains: plasmid-free (blue), carrying the full pOXA-48 plasmid (Tc_MMC-K-0255_J53; red), or carrying a naturally occurring pOXA-48 deletion variant lacking the DUF1496 protein-encoding gene and adjacent sequence (Tf_MMC-K-0151_J53; orange). **b**, Distribution of the DUF1496 protein-encoding gene among *bla*OXA-48- and *bla*OXA-162-harbouring genome assemblies from the NCBI Pathogen Detection database (n = 10,626). Pie charts show, for each geographic region (Africa, the Americas, Asia, Europe, Oceania, and assemblies without geographic metadata), the proportion of assemblies positive (dark blue) or negative (grey) for the gene, with regional assembly counts indicated beside each chart.

### Geographic conservation of the pOXA-48 proteins bounds the approach’s applicability

Because the classification model detects proteins of the pOXA-48 plasmid rather than OXA-48/-162 itself, its diagnostic reach is bounded by the epidemiological distribution of the target element, though non-pOXA-48 carriage represents a minority of OXA-48/-162-producing *Enterobacterales* (see below). To empirically define this boundary, we evaluated the LightGBM-reduced model on isolates carrying *bla*_OXA-48_ on non-IncL plasmid backbones or integrated into the chromosome, together with species-matched negative controls. Sensitivity was reduced to 0.455, while specificity remained high at 0.865, confirming that the spectral signature is specific to the pOXA-48 plasmid context and that the model does not misclassify non-carriers (Extended Data Table 5). The residual signal above zero is consistent with mobile elements such as Tn*6237* carrying a partial pOXA-48 sequence into the chromosome or other plasmid groups, so a fraction of the pOXA-48-encoded protein signature can still be expressed.

The dominant DUF1496 protein is globally highly conserved among pOXA-48 plasmids. Among the IncL-positive subset of these assemblies (6,460 of 10,626), 96.7% carried the DUF1496 protein-encoding gene at a conservative threshold of 100% identity. This indicates that the spectral marker the model relies on is present across essentially the full genetic and geographic range of pOXA-48. Consistent with IncL being the predominant backbone for *bla*_OXA-48/-162_ worldwide, the DUF1496 marker is genomically present in the large majority (70.9%) of *bla*_OXA-48/-162_-carrying isolates overall (Fig. 6b). However, this distribution is uneven and DUF1496 is present in only a minority of assemblies in the Americas (37%) and Oceania (28%) (Fig. 6b), which should be considered for the implementation of this test.

Conversely, the DUF1496 protein is highly specific to the pOXA-48 plasmid context. Among all publicly available complete genomes encoding the exact DUF1496 protein, 98.5% also carried *bla*_OXA-48/-162_ or an OXA-48-like gene. The six remaining records each resided on the conserved pOXA-48 (or a closely related IncL/M) plasmid backbone that did not carry *bla*_OXA-48/-162_. The DUF1496 marker, therefore, barely occurs outside the pOXA-48 plasmid lineage, indicating that detection of the spectral signature is unlikely to generate pOXA-48-unrelated false positives.

## Discussion

Predicting antimicrobial resistance from routine MALDI-TOF mass spectra has been pursued for over a decade across diverse clinically relevant pathogens (reviewed in ^20^). Yet most approaches are trained using phenotypic susceptibility labels, even though the same resistance phenotype can arise through genetically distinct mechanisms. Because MALDI-TOF spectra contain strong phylogenetic signals ^21–23^, models trained on phenotypic endpoints risk capturing lineage-associated correlations rather than the underlying resistance determinants, limiting their transferability across sites, time, and bacterial populations ^24^. This is not unique to phenotype-based models: the commercial Clover MS classifier, which predicts carbapenemase carriage directly and reported 100% sensitivity in its development cohort ^17^, essentially failed to detect our pOXA-48-positive spectra. Because that model treats carbapenemase carriage as a single class spanning structurally unrelated enzymes (OXA-48, KPC, NDM, VIM) on distinct plasmids, it has no common proteomic basis, so its reported accuracy most likely reflected cohort-specific lineage correlations. We therefore reframed the prediction task from inferring a resistance phenotype or genotype to detecting a conserved plasmid through its expressed proteomic signature, with pOXA-48 representing the cleanest case as OXA-48/-162 dissemination is driven predominantly by this highly conserved IncL plasmid rather than by diverse genetic backgrounds ^11^. The closest precedent is the work of Lau et al., who identified KPC-producing *K. pneumoniae* lineages via a MALDI-TOF MS peak of pKpQIL plasmid-associated protein (p019, ∼11,109 Da) ^16^. However, its sensitivity proved dependent on the prevalence of p019-carrying plasmid lineages in a given setting, reaching near-perfect values in endemic pKpQIL-dominated populations like the *K. pneumoniae* ST258 lineage ^25,26^, but failing where *bla*_KPC_ circulates on alternative backbones^27,28^. Whereas they linked one peak to a successful KPC plasmid lineage, we establish a complete mechanistic chain linking a globally conserved plasmid backbone to its encoded proteins, the resulting spectral features, and classifier performance across instruments and species.

Preserved performance after reduction to fewer than 1% of the original input features indicates that the model relies on a compact set of biologically meaningful signals directly linked to pOXA-48, not clonal relatedness or broad spectral similarity. Notably, this performance was achieved with a cohort that is small by machine-learning standards: 246 in-domain isolates versus the >300,000 spectra of phenotypically annotated resources such as DRIAMS ^24^. That a far smaller dataset supports robust, cross-instrument and cross-species generalization underscores that the choice of a biologically meaningful target, a conserved plasmid rather than a phenotype, can be more decisive for model performance than dataset size alone. Targeting the plasmid rather than the host can also explain why the model generalizes to species absent from the training set. For a classifier that exploited host-lineage features, performance on an unseen species would be expected to be around AUROC ≈ 0.5. Instead, the model reached a performance of up to AUROC 0.990 for some unseen species. Such a model transfer to novel host species is, to our knowledge, not achievable with phenotype-based MALDI-TOF models, though validation in rare species remains limited by small sample numbers.

Central to this framework is the DUF1496 domain-containing protein, whose processed mass coincides with the dominant m/z 10,163 feature and whose loss in a natural truncation variant ablates that peak, establishing a direct link from plasmid sequence to spectral signal. Yet no single feature sufficed: in the densely populated whole-cell mass range ^29^, a lone low-abundance peak is easily obscured by chromosomal background and run-to-run variation, whereas the multi-peak signature can stay discriminative when any one feature fails. The approach has nonetheless defined epidemiological boundaries: *bla*_OXA-48_ chromosomal integrations mediated by Tn*6237* ^30^, alternative plasmid backbones such as IncFII, and OXA-48-like variants whose genes circulate on unrelated mobile elements (e.g. *bla*_OXA-232_ on ColKP3 plasmids) fall outside its reach by design. Importantly, these limitations are mechanistically defined and therefore predictable. Unlike phenotype-based classifiers, whose performance can drift as resistance mechanisms and clonal population structures change over time, as in the up-to-0.25 AUROC declines observed within eighteen months of DRIAMS-model deployment ^31^, this framework is constrained primarily by the stability and prevalence of a specific plasmid type. Beyond these biologically defined boundaries, empirical validation was itself limited: training drew on isolates from Germany, Switzerland, and Israel, and external validation spanned two platforms but only two German centres. Furthermore, all spectra came from characterized collections under standardized conditions rather than routine workflows. Prospective multi-centre evaluation addressing both the geographic and the acquisition dimensions is therefore the natural next step toward implementation, and lies beyond the scope of the present study.

Our framework could offer a practical advantage within existing clinical microbiology workflows. Because spectra are acquired routinely for species identification anyway, resistance-associated information can be extracted concurrently, at the earliest diagnostic time point and before susceptibility results are available. The discriminative signals, foremost the DUF1496 marker at m/z 10,163, fall within the default 2–20 kDa acquisition window, so no additional instrumentation, sample preparation, or hands-on time is required. The test is intended not as a standalone diagnostic but as an early screening layer: as universal carbapenemase testing may be constrained by costs or availability, in the case of OXA-48/-162, the model would concentrate rapid confirmatory testing on the isolates most likely to carry pOXA-48, making earlier intervention operationally feasible. Earlier recognition of carbapenemase-producing *Enterobacterales* can prompt pre-emptive contact precautions and confirmatory testing for flagged carriers, which are associated with better infection prevention and control responses ^32^. Furthermore, knowledge of the carbapenemase class is crucial for the appropriate choice of last-resort antibiotics, such as ceftazidime-avibactam for OXA-48/-162 ^33^, and, therefore, earlier detection can prevent inadequate therapy, potentially reducing adverse clinical outcomes ^34^.

More broadly, our results establish a framework for biology-driven MALDI-TOF MS diagnostics in which conserved plasmids are detected through their expressed proteomic signatures. The same principle could extend to other determinants whose dissemination is dominated by conserved plasmid lineages rather than by diverse genetic backgrounds. Strong candidates include *bla*_OXA-181_ on near-identical IncX3 plasmids ^35^, *bla*_NDM-5_ on the stable IncX3 backbone ^36^, and the colistin-resistance determinant *mcr-1* on IncX4 ^37^. With *bla*_NDM_ and *bla*_OXA-48_-like, this list already includes two of the three dominant carbapenemases in European surveillance ^38^. The corollary is that the target must be defined at the determinant–plasmid pair level rather than the resistance-gene family: determinants that move between unrelated plasmids, transposons and chromosomal sites, with *bla*_KPC_ being the clearest example, yield a diluted proteomic signature and would need regionally trained or subtype-resolved models. Operationally, multiple determinant-specific classifiers can be run in parallel on the same spectrum at inference, so broadening coverage is limited only by the availability of a separately validated model per target.

By moving resistance prediction from empirical pattern recognition and database-driven statistical inference toward biologically explainable and mechanism-grounded detection, this framework offers a route to scalable and transferable resistance screening from spectra that clinical laboratories already generate.

## Online methods

### Isolate collections and genomic characterization

Clinical Enterobacterales isolates were collected from routine diagnostic and screening cultures at four institutions for model development (University Hospital Cologne, University of Zurich, Cologne Merheim Medical Centre, and Hadassah Medical Center Jerusalem; collectively termed internal isolates) and two institutions for independent validation (Technical University Munich and University Hospital Frankfurt). Isolates were collected between 2013 and 2025. Negative controls were species-matched to positive isolates by convenience sampling at each site. All isolates were non-duplicate clinical isolates stored as frozen stock at the respective institutions. Sample composition is detailed in Extended Data Table 6.

All isolates were characterized by whole-genome sequencing by either Illumina short read sequencing or Oxford Nanopore long read sequencing at the respective institutions. Sequencing platforms and library preparation methods are detailed in Supplementary Table 2. For short reads, reads were trimmed with Trimmomatic v0.39 and assembled using SPAdes v3.14.1. For long reads, reads were trimmed with Chopper v0.12 and assembled using Flye v2.9.6, polished with Medaka v2.2.2. Resistance genes and plasmid determinants were identified using the ResFinder database and plasmid replicon types using the PlasmidFinder database, applied via ABRicate v1.2.0 (ResFinder and PlasmidFinder databases, snapshot 2025-12-05) with thresholds of 100% identity and 90% coverage. Raw sequence data have been deposited at the Sequence Read Archive (SRA) under BioProject number PRJNA1433510.

The inclusion criterion for the positive class was the genomic co-detection of *bla*_OXA-48/-162_ and an IncL replicon determinant. The negative class comprised isolates lacking any *bla*_OXA-48_-like gene. These definitions applied to all internal and external cohorts.

Within the internal collection, two clinically relevant edge cases were first set aside as separate validation sets to avoid confounding from low-count strata: OXA-48/-162-positive isolates co-harbouring a second carbapenemase gene (double carbapenemases; n = 23 positive, n = 23 species-matched negatives) and OXA-48/-162-positive isolates from species other than *K. pneumoniae* complex, *E. coli*, *Citrobacter freundii* complex, and *E. cloacae* complex (rare species; n = 30 positive, n = 30 species-matched negatives). The main development cohort comprised 153 positives and 153 negatives, the latter comprising 104 carbapenemase-free isolates and 49 carrying non-OXA-48/-162 carbapenemases (predominantly *bla*_KPC_ and *bla*_NDM_), ensuring the classifier discriminates pOXA-48 carriage specifically rather than carbapenemase production in general. This cohort was divided into an in-domain set (n = 246) measured under a standard acquisition protocol and a cross-domain set (n = 60) measured under five acquisition protocols (see MALDI-TOF MS acquisition protocols), stratified by class and species.

The study was reviewed by the Ethics Commission of the Technical University of Munich (reference 2025-110-S-NP), which raised no objections to the project. The study exclusively analysed anonymized bacterial isolates with no access to patient-identifiable data. Therefore, individual informed consent was not required.

### MALDI-TOF MS acquisition protocols

Isolates were cultured from frozen stock onto Columbia blood agar (Biomerieux, Marcy-l’Étoile, France), and an ertapenem 10 µg disc (BioRad, Hercules, CA, USA) was added. After overnight culture, isolates were subcultured from the inner inhibition zone area, again on blood agar. After overnight culture, the isolates were checked for the presence/absence of OXA-48/-162 using a lateral flow test O.K.N.V.I. RESIST-5 (Coris Bioconcept, Gembloux, Belgium). Isolates were then transferred in triplicate for MALDI-TOF measurement on an MSP 96 polished steel target (Bruker Daltonics, Bremen, Germany) and prepared using 1 µl 70% formic acid and, after drying, 1 µl matrix (HCCA for MALDI Biotyper System, Bruker Daltonics). MALDI-TOF MS measurements were conducted on a MALDI Biotyper Sirius IVD System (Bruker Daltonics) with standard settings for species identification. For the internal cross-domain isolates, on top of the abovementioned protocol, different experimental conditions were applied for robustness testing and data augmentation. 1) Culturing on and measuring from a different blood culture brand (Thermo Fisher Scientific, Waltham, MA, USA). 2) Culturing on and measuring from Mueller-Hinton agar (Biomerieux). 3) No use of ertapenem disc on initial culturing (but still assuring OXA-48/-162 presence through lateral flow test). 4) No formic acid in MALDI-TOF measuring.

### Spectral preprocessing and feature engineering

Raw spectra were exported as Bruker FID files and preprocessed using MALDIquant v1.22.3 in R v4.3.3. Intensity values were variance-stabilized by square-root transformation, smoothed using a Savitzky–Golay filter (half-window size 10), baseline-corrected using the SNIP algorithm (iterations = 20 and a second pass with iterations = 100), normalized to total ion current (TIC), and trimmed to the 2,000–20,000 m/z range. Spectra were aligned on the m/z axis by species-specific warping. For each of the four target species, a reference peak list was generated from the in-domain training spectra using MALDIquant’s detectPeaks (signal-to-noise ratio ≥ 2, noise estimation by median absolute deviation, half-window size 20) and referencePeaks (method = “strict”, retaining only peaks present across all spectra within each species group). Warping functions were computed using determineWarpingFunctions with the species-matched reference peak list and applied via warpMassSpectra. External and cross-domain spectra were warped using the same training-derived reference peak lists. Warped spectra were binned into 6,000 bins of 3 Da width spanning the 2,000–20,000 m/z range, with intensities summed within each bin. Four binary features encoding species identity (*K. pneumoniae* complex, *E. coli*, *C. freundii* complex, *E. cloacae* complex) were appended as one-hot-encoded variables, yielding 6,004 input features per spectrum. All features were standardized (zero mean, unit variance) using a scaler fitted exclusively on in-domain training spectra and applied without refitting to all validation and external datasets.

### Commercial baseline (Clover MS)

To establish a commercial baseline, we evaluated Clover MS Data Analysis Software (Clover Bioanalytical Software S.L., Granada, Spain; accessed September 2025), which, to our knowledge, represents the only commercially available system for detecting OXA-48 carbapenemases from MALDI-TOF MS spectra. As the Clover carbapenemase classifier is currently validated only for *K. pneumoniae*, this analysis was restricted to *K. pneumoniae* complex isolates from the internal in-domain and cross-domain datasets (n = 166; 83 OXA-48/-162-positive, 83 OXA-48-negative). Analysis followed the vendor-specified two-stage pipeline as described in Gato et al.^17^ and confirmed by the developer (personal communication). Raw spectra acquired on the MALDI Biotyper Sirius were preprocessed using the vendor’s built-in “Applying Prediction Model Parameters” noise reduction setting, and spectra were analysed with the random-forest binary model reported as best-performing in Gato et al.^17^ for the detection of any carbapenemase. Using the three technical replicates per isolate, carbapenemase positivity was assigned by majority voting (≥2 of 3 replicates positive). For isolates classified as carbapenemase-positive, the same spectra were subsequently passed to the multiclass random-forest subtype module for carbapenemase family assignment (KPC, NDM, OXA-48, VIM). Confidence intervals for sensitivity and specificity were calculated using the Clopper-Pearson exact method.

### Evaluation strategy and cross-validation framework

The 246 in-domain isolates (standard protocol) and 60 cross-domain isolates (measured under all five acquisition protocols) were split using a consistent stratification scheme across all experiments. Cross-domain isolates were divided 50/50 into training and validation sets (approximately 15 positive and 15 negative per split), stratified by class and species, across 20 pre-fixed random seeds. In-domain isolates were assigned exclusively to the training set throughout. All replicates from the same isolate were kept within the same split to prevent data leakage. For evaluation, 100 Monte Carlo iterations were performed per split, randomly selecting one replicate per isolate in each iteration to match deployment conditions. Performance metrics were computed per acquisition protocol per Monte Carlo draw and then averaged, first across Monte Carlo iterations within each protocol, and then across the 20 splits. To increase robustness against temporal batch effects, 50% of training spectra for OXA-48/-162-positive isolates were replaced with spectra from independent measurements of the same isolates acquired approximately one year prior. Because matched re-measurements were not available for negative controls, the corresponding proportion of negative training spectra was drawn from an independently assembled archival collection of species-matched isolates lacking *bla*_OXA-48_-like genes.

Six model configurations were evaluated using these identical splits:

a. Single-protocol baseline (LR): LASSO logistic regression trained on standard-protocol spectra only, evaluated on standard-protocol cross-domain validation isolates and, without retraining, on the four alternative acquisition protocols.
b. Temporal augmentation (LR): As (a), with training data supplemented by archival spectra as described above.
c. Mixed-protocol (LR): Logistic regression trained on standard-protocol spectra combined with cross-domain training isolates across all five protocols, using inverse-proportional condition weighting.
d. Mixed-protocol + temporal (LR): Combination of (b) and (c).
e. Mixed-protocol + temporal (LightGBM): As (d), replacing logistic regression with gradient-boosted decision trees (LightGBM).
f. Leave-one-domain-out (LightGBM): Setup (e) repeated with each protocol sequentially excluded from training and used exclusively for evaluation.

### Model training and hyperparameter settings

Machine-learning models were implemented in Python 3.12.5 using LightGBM v4.5.0 and scikit-learn v1.5.2. Two supervised learning algorithms were evaluated: LASSO logistic regression, as a simple linear classifier, and gradient-boosted decision trees. All models used fixed, a priori hyperparameter settings without hyperparameter optimization. Logistic regression models employed L1 regularization with a regularization strength of C = 0.1, optimized using the SAGA algorithm with a maximum of 1,000 iterations. LightGBM models were trained with the GBDT boosting type, a learning rate of 0.1, and 200 boosting iterations. These values were fixed a priori from common defaults and conventional settings for each algorithm and were deliberately not tuned as a considered choice given the small-sample, high-dimensional nature of the data, to avoid overfitting. Robustness to these choices was confirmed by the sensitivity analysis below (three alternative LightGBM configurations; Supplementary Table 1). Binary classification predictions were obtained at a decision threshold of 0.5 on the predicted probability.

In mixed-protocol training configurations, each spectrum was assigned a weight inversely proportional to the number of training spectra in its acquisition condition: w_i = N / (K × n_k), where N is the total number of training spectra, K is the number of distinct acquisition conditions, and n_k is the number of spectra in the condition to which spectrum i belongs. Weights were normalized to a mean of 1.0 and passed as sample weights during model fitting. This ensures each acquisition condition contributes equally to the loss function despite differing sample sizes.

To assess robustness to hyperparameter choices, three additional LightGBM configurations were evaluated using the same 20-split evaluation framework: reduced model capacity (n_estimators = 100), increased leaf regularization (min_data_in_leaf = 50), and stochastic regularization (feature_fraction = 0.8, bagging_fraction = 0.8, bagging_freq = 1). The untuned baseline configuration was retained for all subsequent analyses.

Performance was assessed using the area under the receiver operating characteristic curve (AUROC), the area under the precision–recall curve (AUPRC), and sensitivity and specificity at a fixed decision threshold of 0.5. AUROC served as the primary metric, as it summarizes ranking performance independently of threshold choice and is robust to the moderate class imbalance in the cohort.

### External evaluation

After cross-validated performance assessment, a final model was trained on all internal data (in-domain training isolates and all cross-domain isolates from both train and validation splits) using the LightGBM configuration and condition weighting scheme described above, and applied without further tuning to independent external datasets. The reduced 58-feature set was selected exclusively on internal validation data prior to external evaluation. 95% confidence intervals for all external rate metrics (AUROC, AUPRC, sensitivity, specificity) were obtained by isolate-level percentile bootstrap (10,000 resamples, stratified by class) on the per-replicate prediction scores, preserving the 100-iteration Monte Carlo replicate-selection used for the point estimate. The two sources of randomness were nested: each outer bootstrap iteration drew one class-stratified isolate resample and, within it, the 100 Monte Carlo replicate selections, and the per-iteration metric was the mean over those 100 selections. Pairwise differences between models were tested with a paired bootstrap that applied the identical isolate resample and replicate selections to every model within each iteration, so that Δ AUROC was evaluated on matched draws and the shared sampling variability cancelled; two-sided P values are the percentile of Δ AUROC.

The Munich dataset comprised 42 clinical isolates (20 OXA-48/-162-positive, 22 OXA-48/-162-negative) acquired on a Bruker Biotyper Sirius one IVD. Raw spectra in Bruker FID format were preprocessed, warped, binned, and standardized identically to the internal data, using the training-derived reference peak lists and scaler. Isolates were collected at the Institute for Medical Microbiology, Immunology, and Hygiene, Technical University of Munich, between 2022 and 2025, and processed using the standard acquisition protocol (ertapenem pre-selection, formic acid overlay, blood agar; see MALDI-TOF MS acquisition protocols).

The Frankfurt dataset comprised 89 clinical isolates (37 OXA-48/-162-positive, 52 OXA-48/-162-negative) acquired on a BioMérieux VITEK MS PRIME system. Raw spectra were exported in mzML format. Averaged spectra were imported using MALDIquantForeign’s importMzMl function and converted to the same tab-separated text format (mass, intensity) used for Bruker data. From this common representation, spectra were preprocessed, warped, binned, and standardized identically to the internal data, using the training-derived reference peak lists and scaler. Applying the Bruker-derived reference peak lists to the VITEK MS spectra was deliberate to have a common reference frame that corrects systematic inter-instrument calibration offsets and places equivalent protein signals in the same m/z bins, which is what allows the internal model to be applied unchanged across platforms.

### Permutation feature importance

To obtain a post-hoc interpretation of the final mixed-protocol LightGBM model, permutation feature importance was computed on internal cross-domain validation data. For each of the 20 splits, feature importance was defined as the decrease in AUROC after random permutation of individual input features, estimated using 30 permutations per feature (scikit-learn permutation_importance). Mean importance values and standard deviations were calculated across permutations and splits, and features were ranked accordingly.

### Feature reduction

To determine the minimal feature set for robust classification, sequential forward feature addition was performed using the permutation importance ranking. Starting from the single most important feature, features were added incrementally in order of decreasing importance. At each step, the LightGBM model was retrained and evaluated using the same 20-split Monte Carlo framework on internal cross-domain data. A coarse screen of feature counts (1, 2, 5, 10, 20, 50, 100, 200, 500, 1,000, 2,000, and all 6,004 features) identified the performance-relevant range, followed by fine-grained evaluation at every integer feature count from 1 to 100. The optimal feature count was selected using the one-standard-error rule: the smallest feature set whose mean AUROC fell within one standard error of the maximum observed AUROC. The reduced model was subsequently evaluated on the external datasets without further tuning.

### Single-feature classification approaches

To assess whether the dominant spectral feature alone could account for classification performance, two complementary single-feature approaches were evaluated, progressing from a machine learning model operating on binned spectra to a rule-based detector operating on unbinned spectra. For the single-feature LightGBM model, the LightGBM model was trained using exclusively the top-ranked feature from the permutation importance analysis (the 3 Da bin centred at m/z 10,163). Training, condition weighting, and evaluation followed the same 20-split Monte Carlo framework as the multi-feature models, with identical hyperparameters. A final single-feature model was trained on all internal data and evaluated on the external Frankfurt dataset without further tuning. For the single-peak detection classifier, a peak detection classifier was evaluated on warped, unbinned spectra using MALDIquant. The classifier assigned OXA-48/-162-positive status if a peak was detected within a specified mass tolerance of m/z 10,163, and negative otherwise. Peak detection parameters were selected by grid search over 128 combinations (noise estimation: MAD, SuperSmoother; half-window size: 3, 5, 10, 20; SNR threshold: 0.25, 0.5, 1, 2; mass tolerance: 2, 4, 6, 8 Da), optimizing AUROC on the full internal training data. The best configuration (SuperSmoother, half-window size 5, SNR 0.25, tolerance 8 Da) was applied to the external Frankfurt dataset using 100 Monte Carlo iterations, each randomly selecting one replicate per isolate, matching the evaluation procedure used for the machine learning models on external data.

### Conjugation and plasmid curing experiments

To test whether the pOXA-48 plasmid is causally responsible for the classifier’s spectral signature, isogenic strain pairs differing exclusively in plasmid content were generated through horizontal gene transfer and plasmid curing. For gain-of-function experiments, pOXA-48 was transferred from the clinical donor strain MMC-K-0255 into two plasmid-free recipients: sodium azide-resistant *E. coli* J53^39^ and *K. quasipneumoniae* PRZ^18^. Liquid mating was performed for 2 h as described previously^14,40^. Transconjugants were selected on coliform chromogenic agar (CCA; Carl Roth, Karlsruhe, Germany) supplemented with 20 µg/mL amoxicillin-clavulanate (Hexal, Holzkirchen, Germany) and 100 µg/mL sodium azide. For loss-of-function, plasmid curing was performed on MMC-K-0225 (*E. coli*). Plasmid curing was achieved with serial passage at elevated temperature (40°C) in antibiotic-free medium for 2 days. Loss of pOXA-48 was confirmed by antibiotic susceptibility testing and whole genome sequencing. Spectra were preprocessed identically to the training data and classified using the LightGBM-reduced model.

### Bottom-up proteomics

#### Isolate selection

Label-free bottom-up proteomics was performed on 17 clinical isolates, eight pOXA-48-positive and nine pOXA-48-negative, spanning *K. pneumoniae*, *E. coli,* and the *E. cloacae* complex. Positive and negative isolates were matched by species and, where feasible, by multilocus sequence type, to control for chromosomal background. When no isolate of the same sequence type was available, the nearest sequence type available was chosen (Supplementary Table 3).

#### Sample preparation

Bacterial isolates were cultured from frozen stock on Columbia blood agar (37 °C, overnight). Colonies (∼1 × 10⁹ cells) were pelleted (10,000 × g, 5 min, 4 °C) and lysed using the TFA-SPEED protocol ^41^. Briefly, pellets were resuspended in 50 µL neat trifluoroacetic acid, vortexed for 30 s, and incubated at 55 °C for 5 min. Lysates were neutralized by the addition of 450 µL 2 M Tris base (final pH 8.2). Protein concentration was determined by tryptophan fluorescence (excitation 280 nm, emission 350 nm). Aliquots containing 20 µg total protein were reduced with 9 mM TCEP and alkylated with 40 mM chloroacetamide (95 °C, 5 min), diluted to 400 µL with ultrapure water, and digested with sequencing-grade trypsin (1:50 enzyme-to-protein ratio, 37 °C, 16 h). Digestion was quenched by acidification to 3% (v/v) formic acid. Peptides were desalted on C18 StageTips (two layers of Empore C18 discs), eluted with 40% acetonitrile/0.1% formic acid, dried by vacuum centrifugation, and reconstituted in 0.1% formic acid for loading onto Evosep tips.

#### Liquid chromatography–tandem mass spectrometry

Peptides were separated on an Evosep Eno system (Evosep, Odense, Denmark) using the 14-samples-per-day method (22.4-min gradient) and analysed on an Orbitrap Astral mass spectrometer (Thermo Fisher Scientific, Waltham, MA, USA, software v2.0-SP1) equipped with a FAIMS Pro interface (compensation voltage −40 V; spray voltage 1,900 V). Data were acquired in data-independent acquisition (DIA) mode: full MS scans in the Orbitrap (m/z 380–980; 120000) were followed by MS2 scans in the Astral analyser across 300 fixed-width 2.0-m/z isolation windows spanning m/z 380–981 (normalized collision energy 25%, 7-ms injection time per window, scan range m/z 150–2,000).

#### Protein sequence database and reference plasmid

A reference pOXA-48 plasmid (RefSeq accession NZ_CP124810.1) carries 83 PGAP-annotated CDS: 81 intact protein-coding CDS, used as the pOXA-48 protein reference, and 2 pseudogenes with partial coordinates. The 2 pseudogenes, together with 2 IS1-family transposase copies encoded across overlapping reading frames by programmed ribosomal frameshifting, are not rendered on the circular plasmid map (Fig. 5b), which therefore shows the remaining 79 CDS. Spectra were searched against strain-level protein FASTA files downloaded from NCBI (*K. pneumoniae:* NC_016845.1, *E. coli*: NC_000913.3, *E. cloacae*: OW968328.1, hereafter the “strain-level FASTA”, which is the database referenced in the Extended Data Fig. 2 legend and reused for the chromosomal differential-abundance analysis).

#### MS-raw file database search

DIA-NN version 2.2.0 was run on a high-performance computing (HPC) cluster to perform parallel searches of all raw files. The library-based search applied a spectral library previously generated from the accumulated strain level FASTA files, as well as the reference pOXA-48 plasmid (total of 33,780 protein entries; 1,531,355 precursors). The library was generated with DIA-NN version 2.2.0 through in silico tryptic digestion, allowing a maximum of one missed cleavage. Modifications included cysteine carbamidomethylation as a fixed modification and both methionine oxidation and N-terminal acetylation as variable modifications, with only one variable modification permitted per peptide. Peptide lengths were set to range from 7 to 30 amino acids. The search parameters were configured with an MS1 mass accuracy of 15 ppm, an MS2 mass accuracy of 15 ppm, and a scan window radius of 6. Match-between-runs was enabled to enhance identification consistency across samples. All other settings were maintained at default values.

#### Protein grouping, quantification, and visualization

From the DIA-NN report, precursor identifications were retained only when all four q-value fields fell below 0.001 — Q.Value < 0.001, PG.Q.Value < 0.001, Lib.Q.Value < 0.001, Lib.PG.Q.Value < 0.001 — corresponding to 0.1% FDR at the precursor, protein-group, and library levels. Filtered identifications were pivoted into a Protein. Group × Run matrix of MaxLFQ intensities and an independent Precursor.Id × Run matrix of normalized precursor intensities. These matrices were restricted to the 17 isolates (8 pOXA-48-positive, 9 pOXA-48-negative) ^42^. DIA-NN’s protein grouping was used unchanged: each Protein.Group is the maximal set of identifiers sharing every quantified peptide, with the leading identifier reported first. A protein group was defined as “pOXA-48-specific” when every semicolon-delimited identifier mapped to the 81 pOXA-48 intact-CDS reference set. Of 59 protein groups whose leading identifier originated from pOXA-48, 58 satisfied this criterion. One mixed group containing one pOXA-48 identifier together with an *Enterobacterales* chromosomal identifier was excluded from all downstream pOXA-48 analyses to avoid attributing chromosomal signal to the plasmid. These 58 groups encompassed 597 precursors used for Fig. 5a.

For the precursor-level heatmap (Fig. 5a), the DIA-NN-reported Precursor.Normalised values (RT-aligned, run-wise normalized by DIA-NN) were log10-transformed without further normalization. Missing values (no quantification at the FDR cascade above) were left as undetected and mapped to the lower bound of the colour scale. Precursors were grouped into the 58 pOXA-48 protein blocks and rows ordered within each block by retention time. Protein blocks themselves were ordered by descending full-length average mass of the leading protein computed with Bio.SeqUtils.ProtParam.ProteinAnalysis (Biopython ^43^), shown by the left colour bar (∼7.5 to ∼125 kDa). Column order was determined hierarchically by pOXA-48 group, then by species, then by total summed log10 intensity within each (group, species) stratum so that the categorical column annotations read as contiguous blocks above the heatmap.

For the circular plasmid map (Fig. 5b), each of the 79 rendered pOXA-48 CDS was drawn as a directional arrow on its strand using pycirclize, with the genomic coordinates taken from the [location=…] field of the species-level FASTA. For each CDS, the per-isolate PG.MaxLFQ of any pOXA-48 protein group containing that identifier was retrieved, restricted to the 8 pOXA-48-positive isolates, and summarized as log10 of the across-sample median. CDSs were coloured by this log10-median intensity (plasma colour map) when at least one pOXA-48-positive isolate quantified the protein group. CDSs not quantified in any pOXA-48-positive isolate were rendered in mid-grey.

Per-isolate precursor coverage (Supplementary Table 4) was tabulated by counting, for each of the 17 isolates, the number of the 597 pOXA-48-specific precursors with a quantified Precursor.Normalised value at the same 0.1% FDR cascade. Per-isolate counts were summarized as the within-class median, and the between-class difference was expressed as the ratio of the positive- to negative-class median count. All transformations, filtering, statistics, and figure generation were implemented in Python 3.11 and rendered with matplotlib and pycirclize.

#### Signal-peptide prediction and maturation-aware molecular weights

Signal peptides on the 58 detected pOXA-48–encoded proteins were predicted with SignalP 6.0 run with the academic distribution (version 6.0i, slow-sequential mode, --organism other) ^44^. In brief, SignalP 6.0 couples a ProtBert protein-language-model encoder to a conditional random-field decoder that labels each residue as belonging to the n-, h-, or c-region of one of five signal-peptide classes — Sec/SPI, Sec/SPII (lipoprotein), Tat/SPI, Tat/SPII, or Sec/SPIII (pilin) — or to the mature chain. The slow-sequential ensemble averages predictions across six independently trained checkpoints. For each protein, we recorded the predicted signal-peptide class, the inferred cleavage-site position, and the associated cleavage probability. Of the 58 detected pOXA-48 proteins, 13 were predicted to carry a signal peptide (10 Sec/SPI and 3 Sec/SPII lipoproteins). No Tat or pilin substrates were detected. Mature sequences were defined as the residues C-terminal to the predicted cleavage site for proteins assigned to any signal-peptide class and as the full chain otherwise. Average molecular weights of the full and mature sequences were computed with Bio.SeqUtils.ProtParam.ProteinAnalysis (Biopython ^43^), after removal of stop (*) and ambiguous (X) residues, and are reported alongside the full-length values in Supplementary Table 5.

#### Differential abundance of chromosomal proteins

To confirm that the class-discriminating signal originates specifically from plasmid-borne gene products, chromosomal proteins were tested for differential abundance between pOXA-48-positive and pOXA-48-negative isolates separately for each species (*K. pneumoniae, E. coli, E. cloacae* complex). For each species, the analysis universe was restricted to protein groups whose every semicolon-delimited identifier mapped to that species’ reference FASTA. Protein groups containing any pOXA-48 identifier were explicitly removed (59 pOXA-48-touching groups per species), so the test reports only chromosomal background. Per-protein-group intensities were log10-transformed, and a Welch’s two-sample t-test was performed between the pOXA-48-positive and pOXA-48-negative runs of that species. Protein groups quantified in fewer than two samples per class were skipped. Per-species P values were adjusted for multiple testing using the Benjamini–Hochberg procedure.

#### Mapping discriminative MALDI-TOF features to pOXA-48 proteins and chance-match control

The predicted average masses of the 58 detected pOXA-48 proteins were cross-referenced against the m/z positions of the 58 retained LightGBM features. A feature was counted as protein-matched when a predicted protein mass fell within ±10 Da of the feature’s m/z-bin centre, considering both singly-charged ([M+H]⁺) and doubly-charged ([M+2H]²⁺) ions. Mature masses were used for proteins with a predicted signal peptide and full-length masses otherwise. Each feature’s representative m/z was defined as the centre of its 3-Da bin (left bin-edge + 1.5 Da). Multiple proteins, or both ion species of the same protein, matching the same feature were counted once per feature rather than multiplicatively. Conversely, a single protein matching several adjacent features was counted as a match for each of those features individually. To assess whether the number of matches exceeded chance expectation, a permutation null distribution was constructed by redistributing the 58 feature positions uniformly at random (without replacement) across the 6,000 m/z bins of the binned spectrum, while the 58 predicted protein masses (yielding 116 candidate ion m/z values: [M+H]⁺ and [M+2H]²⁺) were held fixed, and recounting the number of protein-matched features. This was repeated 10,000 times. The empirical P value was defined as the fraction of permutations yielding at least as many matches as observed (one-sided, using the same matching rule).

### Transformation with naturally truncated pOXA-48

To establish whether the m/z 10,163 spectral feature requires an intact DUF1496 protein open reading frame, the naturally truncated pOXA-48 variant pOXA-48.7c (Extended Data Fig. 3) was transferred into a plasmid-free recipient. Donor isolate MMC-K-0151 (identical to *C. freundii* CF17067 previously characterized by Sattler et al. ^19^, BioSample SAMN25408304) carries pOXA-48.7c (63,126 bp), in which a natural deletion extending from the left inverted repeat of Tn*7442* into the plasmid backbone has removed both the *trb* conjugation locus and the gene encoding the DUF1496 protein, while leaving *bla*_OXA-48_ intact. Because loss of the *trb* operon abolishes mating-pair formation and renders the plasmid non-conjugative ^19^, plasmid DNA was extracted from MMC-K-0151 using the Plasmid Maxi Kit (Qiagen) and introduced into electrocompetent *E. coli* J53 by electroporation, as described previously ^14^. Transformants (Tf_MMC-K-0151_J53) were selected on CCA supplemented with 20 µg/mL amoxicillin-clavulanate, and uptake of the truncated plasmid was confirmed by a lateral flow test for OXA-48/-162 production and by whole-genome sequencing.

### Sensitivity analysis on non-IncL *bla*_OXA-48_ carriers

To define the diagnostic boundary of the pOXA-48-targeted classification model, isolates carrying *bla*_OXA-48/-162_ on non-IncL plasmid backbones or integrated into the chromosome were evaluated as a separate validation set (n = 32 positive, n = 32 species-matched negatives, Extended Data Table 5). Positive isolates were identified from the internal collections based on whole-genome sequencing. Inclusion required detection of *bla*_OXA-48/-162_ by ResFinder in the absence of an IncL replicon on that contig by PlasmidFinder (thresholds as above). Negative controls were species-matched isolates without OXA-48-like carbapenemase. Spectra were acquired, preprocessed, and classified using the LightGBM-reduced model identically to the other validation sets, with performance assessed using 100 Monte Carlo iterations.

### *In silico* conservation and specificity analysis

All available genome assemblies harbouring *bla*_OXA-48_ or *bla*_OXA-162_ were downloaded on 22 January 2026 from NCBI Pathogen Detection (beta) using the query “AMR_genotypes:blaOXA-48” and “AMR_genotypes:blaOXA-162” (total assemblies: *bla*_OXA-48_ = 10,570, *bla*_OXA-162_ = 56). Presence of the DUF1496 protein-encoding gene was assessed with ABRicate v1.2.0 (Seemann T, Abricate, GitHub https://github.com/tseemann/abricate) using a custom-made database, with thresholds set to 100% identity and 90% coverage. IncL-replicon carriage was determined for the same assemblies with the PlasmidFinder database (threshold 100% identity, 90% coverage).

Conversely, to assess whether the DUF1496 protein occurs outside the pOXA-48 context, the full-length DUF1496 protein sequence encoded by the reference pOXA-48 plasmid (RefSeq NZ_CP124810.1) was queried with tblastn (BLAST+ 2.16.0) against the NCBI nucleotide database (nt), restricted to complete bacterial genomic records (accessed 2 June 2026). Records matching at 100% amino-acid identity over the full protein length (100% query coverage) were retained as exact DUF1496 carriers (n = 393). For each carrier replicon, carbapenemase genes and plasmid replicon types were identified with ABRicate v1.2.0 (ResFinder/NCBI and PlasmidFinder databases, snapshot 2025-12-05; 90% identity, 90% coverage), and co-occurrence with *bla*_OXA-48_-like and other carbapenemase genes was recorded.

### Use of large language models

During the preparation of this manuscript, the authors used Claude (Anthropic) to edit and, in places, to draft and rewrite portions of the text for clarity and readability. The analysis code was developed with assistance from Claude Code (Anthropic). All AI-assisted text and code were reviewed, verified, and edited by the authors, who take full responsibility for the content of the manuscript.

## Data availability

The MALDI-TOF mass spectra and associated metadata generated in this study have been deposited at Zenodo (https://doi.org/10.5281/zenodo.19206415) under a CC-BY-4.0 license. Short read sequencing reads for the internal OXA-48/-162-positive isolates, and long reads for the external Frankfurt isolates are available at the NCBI Sequence Read Archive under BioProject PRJNA1433510. Long reads for the external Munich cohort are available under BioProject PRJNA1297122 (MMC-M-0015 = SRR34727958, MMC-M-0016 = -953, MMC-M-0017 = -950, MMC-M-0018 = -955, MMC-M-0019 = -947, MMC-M-0020 = -948), and raw reads of the remaining isolates at Zenodo (https://zenodo.org/records/20049839). The reference pOXA-48 plasmid (RefSeq NZ_CP124810.1) and the reference genomes used for the proteomic database search (*K. pneumoniae* NC_016845.1, *E. coli* NC_000913.3, *E. cloacae* OW968328.1) are available from NCBI. Publicly available *bla*_OXA-48/OXA-162_ genome assemblies were obtained from NCBI Pathogen Detection (accessed 22 January 2026). Resistance gene and replicon calls used the ResFinder and PlasmidFinder databases (snapshot 2025-12-05).

## Code availability

Custom code for spectral preprocessing, model training, feature-importance and reduction analyses, and the proteomic feature-mapping is available at Zenodo (https://doi.org/10.5281/zenodo.21241413).

## Acknowledgements

We acknowledge Yvonne Stoll, Vanessa Mlottek from the University of Oldenburg, and Diana Albertos Torres, Ana Andrade Barrios, Klara Haldimann, Natalia Kolesnik-Goldmann, and Daniel Gander from the University of Zurich for technical assistance. We thank Silke Peter from the University of Tübingen and the Microbial Genomics Division at the Institute for Medical Microbiology of the University of Zurich for their valuable support with genome sequencing of clinical isolates. We thank Coris Bioconcept (Gembloux, Belgium) for providing the O.K.N.V.I. RESIST-5 lateral flow assays used in this study at a reduced cost. We thank the Clinical Microbiology Laboratory team at Hadassah Medical Center for technical assistance.

## Funding Statements

J.Sa. discloses support for the research of this work from the German Center for Infection Research (DZIF, grant number TTU 08.923). J.So. discloses support for the research of this work from the Deutsche Forschungsgemeinschaft (DFG, German Research Foundation, grant number 493624332). S.G. discloses support for the research of this work from the Dr. Rolf M. Schwiete-Stiftung. All other authors declare no relevant funding.

## Competing interests

K.B. is co-founder and scientific advisor of Computomics GmbH, Tübingen, Germany. A.F.W. has received speaker honoraria from Cepheid. All other authors declare no competing interests.

## Author contributions

- Conceptualization: J.Sa., K.B., A.H., A.E.
- Methodology: J.Sa., K.B., J.M.-R., D.C., L.M., J.M., T.S.
- Software: J.Sa.
- Formal analysis: J.Sa., J.M.-R., P.T., D.C., L.M., J.M.
- Investigation: J.Sa., J.M.-R., P.T., J.So., S.G., T.P., H.M.B.S.-S., T.R.
- Resources: J.So., S.G., J.J., A.F.W., J.M.-G., A.E., A.H., T.P., E.S., M.M.
- Data curation: J.Sa., J.R., Y.G., E.S., H.M.B.S.-S., T.R.
- Visualization: J.Sa.
- Validation: J.Sa.
- Project administration: J.Sa.
- Supervision: K.B., A.H., A.E., M.M.
- Writing – original draft: J.Sa.
- Writing – review & editing: all authors

## Extended data

**Extended Data Table 1.** Commercial-baseline (Clover MS) evaluation on *Klebsiella pneumoniae* complex isolates. Performance of the Clover MS Data Analysis Software binary carbapenemase-detection model on 166 *Klebsiella pneumoniae* complex isolates from the internal datasets (83 OXA-48-positive, 83 OXA-48-negative; Methods). Carbapenemase positivity was assigned per isolate by majority vote across three technical replicates (≥2 of 3 positive). Sensitivity and specificity 95% confidence intervals are Clopper–Pearson exact. The single detected positive isolate received discordant subtype calls (one replicate OXA-48, one KPC). AUROC/AUPRC are not defined because the classifier returns a binary call rather than a probability score. TP, FN, FP, TN, confusion-matrix counts.

| Comparator | Sensitivity | Specificity | TP | FN | FP | TN |
| --- | --- | --- | --- | --- | --- | --- |
| Clover MS (binary carbapenemase detection model) | 0.012 (0.000-0.065) | 1.000 (0.957-1.000) | 1 | 82 | 0 | 83 |

**Extended Data Table 2.** Classification performance across model configurations and acquisition protocols. Performance of six model configurations, each evaluated on the five acquisition protocols (standard; ertapenem pre-selection omitted; formic-acid overlay omitted; Mueller–Hinton agar; alternative blood agar). Configurations: LASSO logistic regression (LR) trained on standard-protocol spectra only; LR with temporal augmentation (training combined across two measurement time points); LR mixed-protocol (standard-protocol plus cross-domain isolates with inverse-proportional condition weighting); LR mixed-protocol + temporal; LightGBM mixed-protocol + temporal; and the leave-one-domain-out (LODO) LightGBM. AUROC, AUPRC (area under the precision–recall curve), sensitivity, and specificity are means (s.d.) across 20 random seeds, with sensitivity and specificity at a 0.5 decision threshold. TP, FN, FP, and TN are mean per-seed counts of true positives, false negatives, false positives, and true negatives.

| Model configuration | Protocol | AUROC (±s.d.) | AUPRC (±s.d.) | Sensitivity (±s.d.) | Specificity (±s.d.) | TP | FN | FP | TN |
| --- | --- | --- | --- | --- | --- | --- | --- | --- | --- |
| LR, single-protocol | Standard | 0.979 (0.011) | 0.980 (0.010) | 0.956 (0.025) | 0.893 (0.029) | 14.5 | 0.7 | 1.6 | 13.3 |
| | $\Delta$ Ertapenem | 0.833 (0.037) | 0.845 (0.035) | 0.619 (0.060) | 0.902 (0.029) | 9.4 | 5.8 | 1.5 | 13.4 |
| | $\Delta$ Formic acid | 0.786 (0.042) | 0.789 (0.047) | 0.550 (0.075) | 0.827 (0.036) | 8.3 | 6.8 | 2.6 | 12.3 |
|  | Mueller-Hinton | 0.803 (0.027) | 0.829 (0.029) | 0.490 (0.066) | 0.903 (0.031) | 7.4 | 7.7 | 1.4 | 13.4 |
|  | Alt. blood agar | 0.787 (0.048) | 0.789 (0.051) | 1.000 (0.000) | 0.188 (0.037) | 15.2 | 0 | 12.1 | 2.8 |
| LR, temporal augmentation | Standard | 0.958 (0.014) | 0.963 (0.013) | 0.934 (0.019) | 0.845 (0.037) | 14.2 | 1 | 2.3 | 12.5 |
| | $\Delta$ Ertapenem | 0.931 (0.016) | 0.926 (0.020) | 0.831 (0.054) | 0.872 (0.042) | 12.6 | 2.6 | 1.9 | 12.9 |
| | $\Delta$ Formic acid | 0.953 (0.015) | 0.954 (0.014) | 0.908 (0.026) | 0.839 (0.045) | 13.8 | 1.4 | 2.4 | 12.5 |
|  | Mueller-Hinton | 0.915 (0.022) | 0.915 (0.025) | 0.652 (0.064) | 0.924 (0.035) | 9.9 | 5.3 | 1.1 | 13.7 |
|  | Alt. blood agar | 0.924 (0.024) | 0.931 (0.023) | 0.784 (0.061) | 0.910 (0.029) | 11.9 | 3.3 | 1.3 | 13.5 |
| LR, mixed-protocol | Standard | 0.947 (0.022) | 0.954 (0.021) | 0.868 (0.045) | 0.874 (0.066) | 13.2 | 2 | 1.9 | 13 |
| | $\Delta$ Ertapenem | 0.911 (0.031) | 0.926 (0.024) | 0.763 (0.083) | 0.893 (0.045) | 11.5 | 3.6 | 1.6 | 13.2 |
| | $\Delta$ Formic acid | 0.950 (0.029) | 0.952 (0.032) | 0.854 (0.068) | 0.907 (0.064) | 12.9 | 2.2 | 1.4 | 13.5 |
|  | Mueller-Hinton | 0.923 (0.038) | 0.930 (0.033) | 0.760 (0.088) | 0.888 (0.043) | 11.5 | 3.6 | 1.7 | 13.2 |
|  | Alt. blood agar | 0.934 (0.027) | 0.938 (0.029) | 0.817 (0.080) | 0.882 (0.042) | 12.4 | 2.8 | 1.8 | 13.1 |
| LR, mixed + temporal | Standard | 0.923 (0.021) | 0.930 (0.023) | 0.816 (0.061) | 0.867 (0.054) | 12.3 | 2.8 | 2 | 12.9 |
|  | Δ Ertapenem | 0.917 (0.025) | 0.923 (0.022) | 0.813 (0.079) | 0.884 (0.040) | 12.3 | 2.8 | 1.7 | 13.1 |
|  | Δ Formic acid | 0.968 (0.019) | 0.972 (0.018) | 0.915 (0.049) | 0.872 (0.054) | 13.8 | 1.3 | 1.9 | 12.9 |
|  | Mueller-Hinton | 0.939 (0.030) | 0.947 (0.024) | 0.820 (0.101) | 0.888 (0.065) | 12.4 | 2.7 | 1.7 | 13.2 |
|  | Alt. blood agar | 0.942 (0.024) | 0.947 (0.025) | 0.849 (0.071) | 0.879 (0.042) | 12.9 | 2.3 | 1.8 | 13 |
| LightGBM, mixed + temporal | Standard | 0.951 (0.015) | 0.947 (0.019) | 0.891 (0.035) | 0.896 (0.034) | 13.5 | 1.7 | 1.6 | 13.3 |
|  | Δ Ertapenem | 0.967 (0.014) | 0.968 (0.014) | 0.891 (0.040) | 0.917 (0.028) | 13.5 | 1.6 | 1.2 | 13.6 |
|  | Δ Formic acid | 0.975 (0.010) | 0.977 (0.011) | 0.921 (0.032) | 0.888 (0.040) | 14 | 1.2 | 1.7 | 13.2 |
|  | Mueller-Hinton | 0.957 (0.011) | 0.958 (0.012) | 0.873 (0.055) | 0.887 (0.044) | 13.2 | 1.9 | 1.7 | 13.2 |
|  | Alt. blood agar | 0.965 (0.013) | 0.967 (0.013) | 0.917 (0.036) | 0.873 (0.034) | 13.9 | 1.3 | 1.9 | 13 |
| LODO, LightGBM, mixed + temporal | Standard | 0.940 (0.017) | 0.935 (0.024) | 0.875 (0.046) | 0.887 (0.027) | 13.3 | 1.9 | 1.7 | 13.2 |
|  | Δ Ertapenem | 0.961 (0.012) | 0.962 (0.013) | 0.885 (0.038) | 0.893 (0.027) | 13.4 | 1.7 | 1.6 | 13.3 |
|  | Δ Formic acid | 0.966 (0.012) | 0.968 (0.013) | 0.932 (0.029) | 0.830 (0.049) | 14.1 | 1 | 2.5 | 12.3 |
|  | Mueller-Hinton | 0.952 (0.014) | 0.954 (0.012) | 0.851 (0.065) | 0.889 (0.040) | 12.9 | 2.3 | 1.6 | 13.2 |
|  | Alt. blood agar | 0.960 (0.017) | 0.962 (0.018) | 0.919 (0.044) | 0.861 (0.032) | 13.9 | 1.2 | 2.1 | 12.8 |

**Extended Data Table 3.** External-cohort evaluation on the Munich and Frankfurt datasets. Performance of four classifiers — LightGBM full (6,004 features), LightGBM reduced (58 features), LightGBM restricted to the top-ranked feature (the m/z 10,163 bin), and a rule-based single-peak detector at m/z 10,163 — trained on internal data and applied without further tuning to the Munich (n = 42) and Frankfurt (n = 89) cohorts. AUROC, AUPRC, sensitivity, and specificity (the latter two at a 0.5 decision threshold) are shown as estimates (95% CI): point estimates are the mean of 100 Monte Carlo spectrum draws on the full external cohort, and 95% confidence intervals are from an isolate-level Monte Carlo-aware percentile bootstrap (10,000 resamples, stratified by class). The confusion-matrix counts (TP, FN, FP, TN) are mean counts over the same 100 draws.

| Metric | AUROC | AUPRC | Sensitivity | Specificity | TP | FN | FP | TN |
| --- | --- | --- | --- | --- | --- | --- | --- | --- |
| **Munich full features** | 0.978 (0.954-0.995) | 0.980 (0.960-0.995) | 0.896 (0.810-0.971) | 0.892 (0.825-0.956) |  | 2.1 | 2.4 | 19.6 |
| **Munich reduced** | 0.982 (0.964-0.996) | 0.983 (0.966-0.996) | 0.931 (0.870-0.985) | 0.880 (0.793-0.953) |  | 1.4 | 2.6 | 19.4 |
| **Munich Top 1** | 0.862 (0.790-0.927) | 0.814 (0.730-0.915) | 0.885 (0.795-0.963) | 0.713 (0.587-0.830) |  | 2.3 | 6.3 | 15.7 |
| **Munich single peak** | 0.891 (0.805-0.956) | 0.873 (0.778-0.955) | 1.000 (1.000-1.000) | 0.241 (0.107-0.389) |  | 0.0 | 16.7 | 5.3 |
| **Frankfurt full features** | 0.951 (0.918-0.979) | 0.936 (0.884-0.974) | 0.844 (0.760-0.924) | 0.949 (0.903-0.986) |  | 5.8 | 2.6 | 49.4 |
| **Frankfurt reduced** | 0.964 (0.937-0.984) | 0.951 (0.911-0.980) | 0.819 (0.733-0.896) | 0.954 (0.916-0.987) |  | 6.7 | 2.4 | 49.6 |
| **Frankfurt Top 1** | 0.809 (0.728-0.879) | 0.709 (0.604-0.806) | 0.485 (0.370-0.585) | 0.834 (0.748-0.909) |  | 19.1 | 8.6 | 43.4 |
| **Frankfurt single peak** | 0.856 (0.802-0.914) | 0.807 (0.715-0.878) | 0.825 (0.734-0.914) | 0.817 (0.755-0.881) |  | 6.5 | 9.5 | 42.5 |

**Extended Data Table 4.** Per-species external performance of the LightGBM-reduced model. The model was evaluated on the Munich (n = 42) and Frankfurt (n = 89) cohorts, stratified by host species. AUROC, AUPRC, sensitivity and specificity (the latter two at a 0.5 threshold) are shown as estimates (95% CI): point estimates are the mean of 100 Monte Carlo spectrum draws (one replicate per isolate per draw), and 95% confidence intervals are from an isolate-level Monte Carlo-aware percentile bootstrap (10,000 resamples, stratified by class). Confusion-matrix counts (TP, FN, FP, TN) are mean counts over the same draws. Per-species rows partition the same draws. n, total isolates per stratum; n.e., interval not estimable (fewer than 5 isolates per class). Per-species estimates with small n should be interpreted with caution.

| **Dataset** | **n isolates** | **AUROC** | **AUPRC** | **Sensitivity** | **Specificity** | **TP** | **FN** | **FP** | **TN** |
| --- | --- | --- | --- | --- | --- | --- | --- | --- | --- |
| **Munich (all species)** | 42 | 0.982 (0.964-0.996) | 0.983 (0.966-0.996) | 0.931 (0.870-0.985) | 0.880 (0.793-0.953) |  | 1.4 | 2.6 | 19.4 |
| --<br>*Citrobacter freundii*<br>complex | 23 | 0.994 (0.980-1.000) | 0.994 (0.982-1.000) | 0.937 (0.853-1.000) | 0.889 (0.795-0.974) |  | 0.7 | 1.3 | 10.7 |
| --<br>*Escherichia coli* | 6 | 1.000 (n.e.) | 1.000 (n.e.) | 1.000 (n.e.) | 1.000 (n.e.) |  | 0 | 0 | 3 |
| --<br>*Klebsiella pneumoniae*<br>complex | 13 | 0.933 (0.855-0.992) | 0.940 (0.876-0.992) | 0.883 (0.757-1.000) | 0.814 (0.616-0.963) |  | 0.7 | 1.3 | 5.7 |
| **Frankfurt**<br>**(all species)** | 89 | 0.964 (0.937-0.984) | 0.951 (0.911-0.980) | 0.819<br>(0.733-0.896) | 0.954<br>(0.916-0.987) |  | 6.7 | 2.4 | 49.6 |
| --<br>*Citrobacter*<br>*freundii*<br>*complex* | 11 | 1.000 (1.000-1.000) | 1.000 (1.000-1.000) | 0.926<br>(0.796-1.000) | 1.000<br>(1.000-1.000) |  | 0.4 | 0 | 6 |
| --<br>*Enterobact*<br>*er cloacae*<br>*complex* | 10 | 0.994 (0.978-1.000) | 0.995 (0.983-1.000) | 0.880<br>(0.714-1.000) | 1.000<br>(1.000-1.000) |  | 0.6 | 0 | 5 |
| --<br>*Escherichia*<br>*coli* | 20 | 0.981 (0.951-1.000) | 0.966 (0.916-1.000) | 0.618<br>(0.318-0.887) | 1.000<br>(1.000-1.000) |  | 2.3 | 0 | 14 |
| --<br>*Klebsiella*<br>*pneumonia*<br>*e complex* | 48 | 0.954 (0.915-0.984) | 0.940 (0.883-0.981) | 0.836<br>(0.740-0.932) | 0.912<br>(0.844-0.974) |  | 3.4 | 2.4 | 24.6 |

**Extended Data Table 5.**
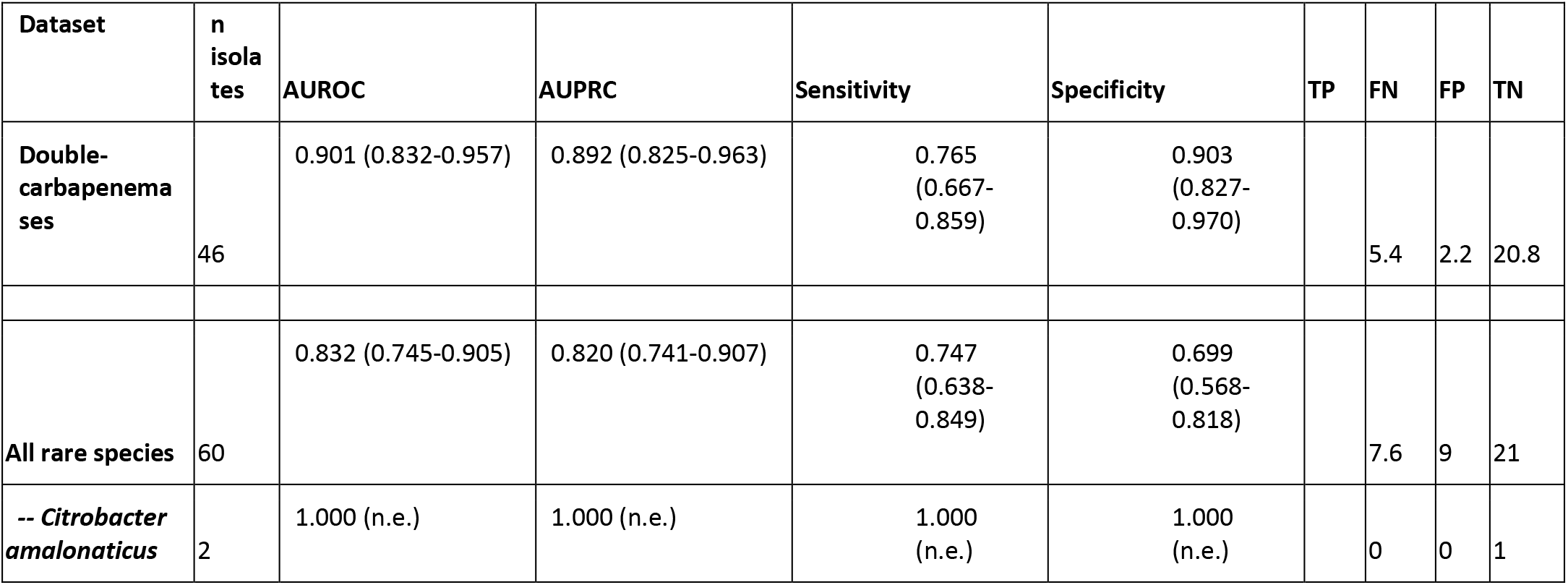
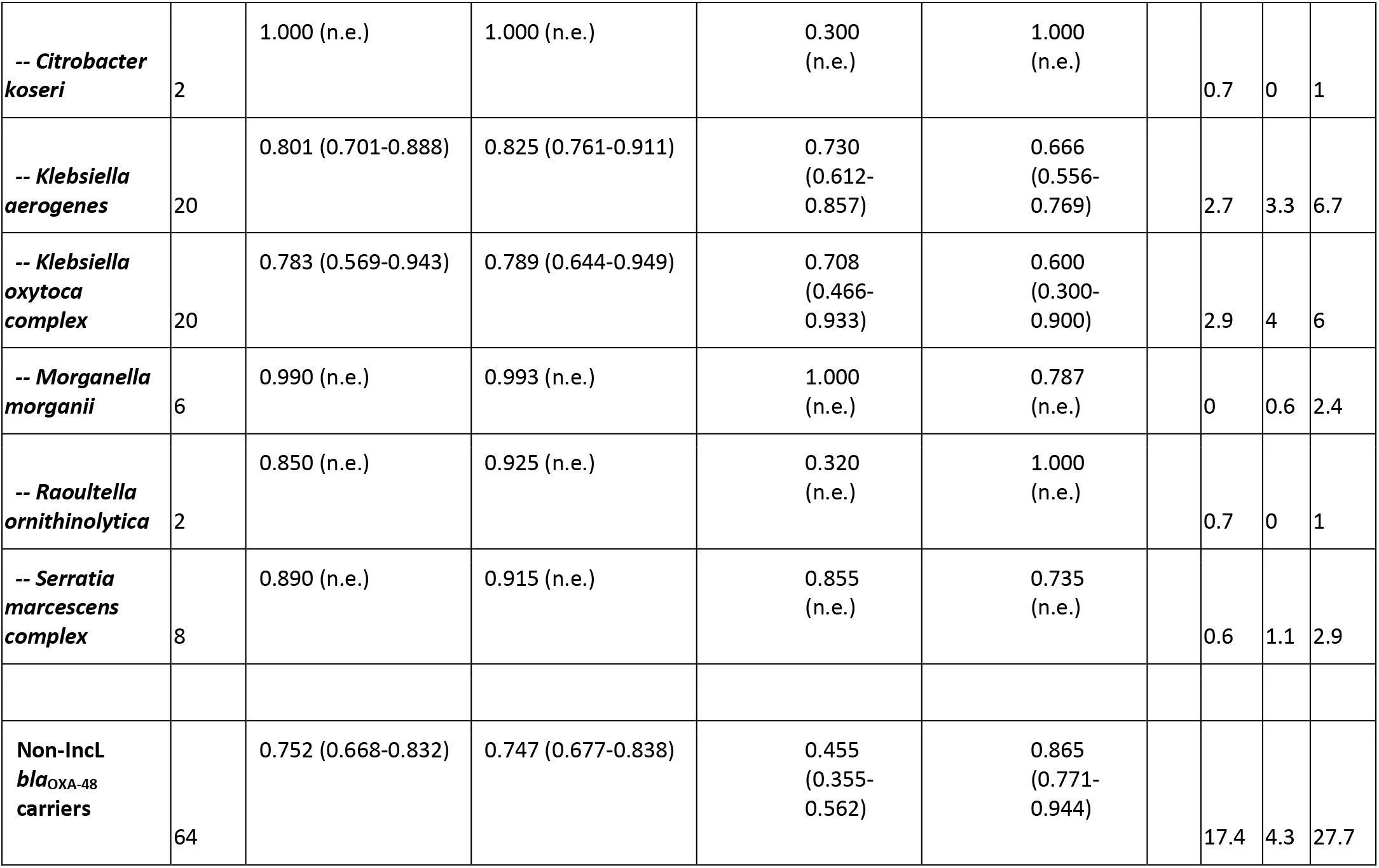
Performance of the LightGBM-reduced model on additional validation subsets. Classification performance of the LightGBM-reduced model on isolate subsets excluded from training: double-carbapenemase producers (n = 46); rare species not represented in training, reported in aggregate (n = 60) and per species; and isolates carrying blaOXA-48 on non-IncL plasmid backbones or the chromosome (n = 64). Columns: number of isolates (n), AUROC, AUPRC, sensitivity and specificity (at a 0.5 decision threshold), and confusion-matrix counts (TP, FN, FP, TN). Derived metrics are shown as estimates (95% CI): point estimates are the mean of 100 Monte Carlo spectrum draws, and 95% confidence intervals are from an isolate-level Monte Carlo-aware percentile bootstrap (10,000 resamples, stratified by class); n.e., interval not estimable (fewer than 5 isolates per class). Confusion-matrix counts are mean counts over the same draws. As per-species isolate numbers are small and uneven (2–20 per species), individual per-species estimates should be interpreted with caution.

**Extended Data Fig. 1.**
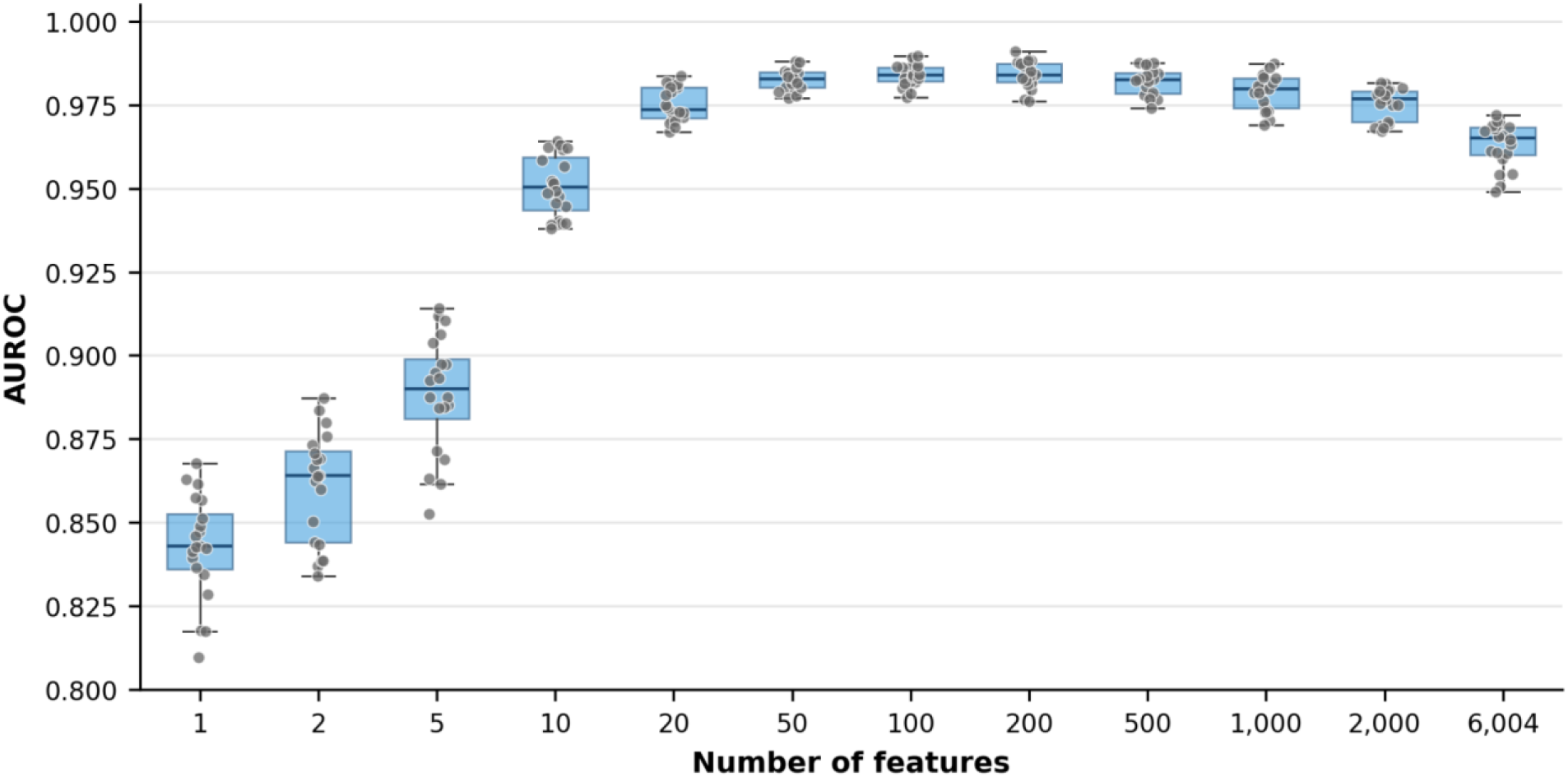
Classification performance peaks at intermediate feature counts. Distribution of AUROC across 20 random seeds for each of twelve feature counts (1, 2, 5, 10, 20, 50, 100, 200, 500, 1,000, 2,000, and all 6,004 features), evaluated with the multi-protocol LightGBM model and averaged across the five acquisition protocols. Box plots show the median (centre line) and interquartile range (box); whiskers extend to the most extreme points within 1.5× the interquartile range; individual seed values (n = 20 per feature count) are overlaid as grey dots.

**Extended Data Fig. 2.**
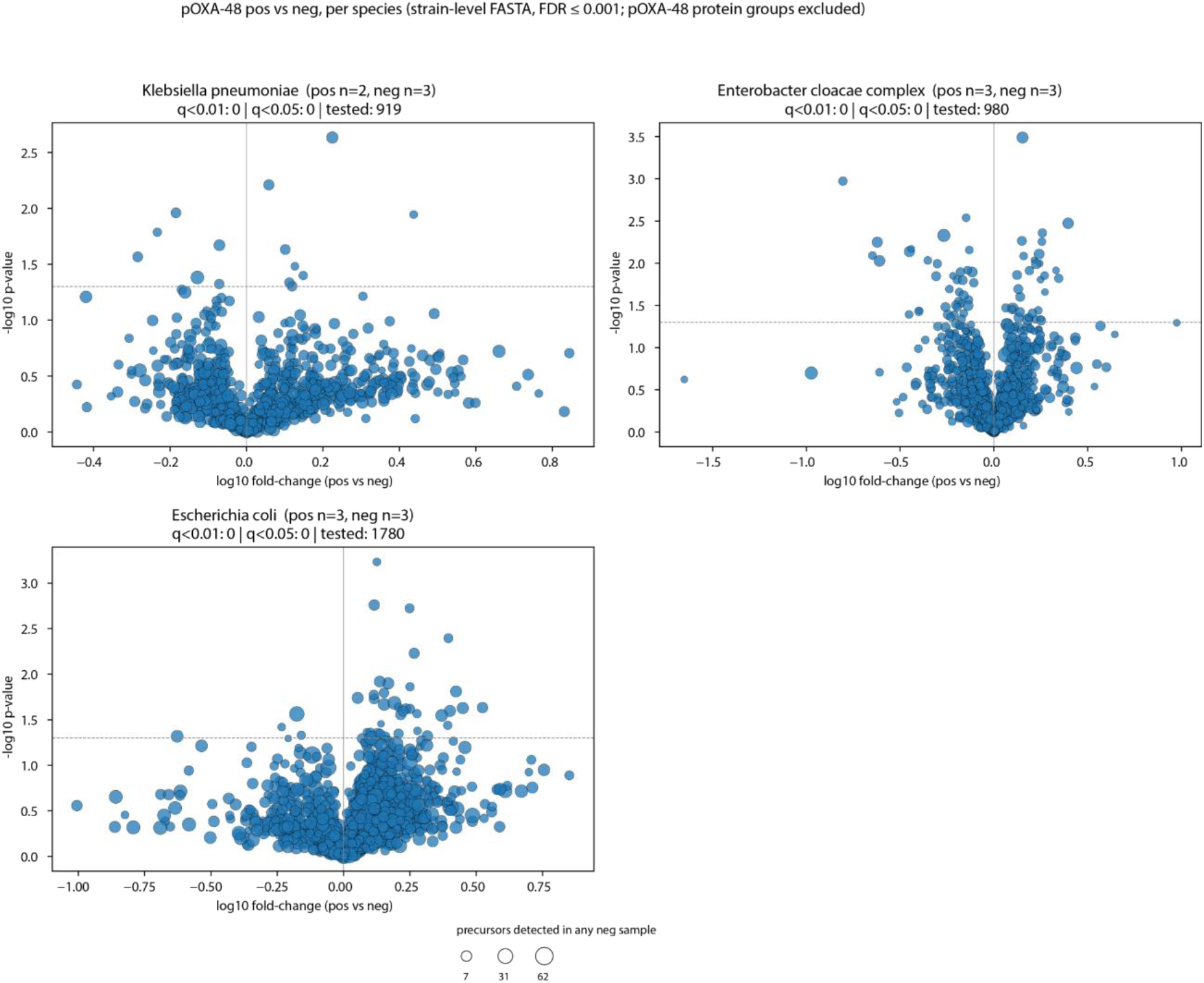
Chromosomal protein abundance does not discriminate pOXA-48-positive from pOXA-48-negative isolates. Volcano plots of differential abundance for chromosomally encoded protein groups between pOXA-48-positive and pOXA-48-negative isolates, shown separately for *Klebsiella pneumoniae* (positive n = 2, negative n = 3), the E*nterobacter cloacae* complex (n = 3, n = 3), and *Escherichia coli* (n = 3, n = 3). For each species, the analysis was restricted to protein groups mapping entirely to that species’ reference proteome, with all pOXA-48-associated protein groups removed. Each point is a protein group: x-axis, log10 fold-change (pOXA-48-positive versus pOXA-48-negative); y-axis, –log10 P value from a Welch’s two-sample t-test on log10-transformed intensities, with per-species Benjamini–Hochberg correction. Point size scales with the number of precursors detected in any negative-class sample (legend). The dashed horizontal line marks P = 0.05. Panel headers give the number of protein groups significant at q < 0.01 and q < 0.05, and the total number tested per species. Precursors were identified at 0.1% FDR.

**Extended Data Table 6.** Species composition and sample sizes across datasets. Number of OXA-48/OXA-162-positive (pos) and negative (neg) isolates per species group for each dataset used in model development and validation (internal in-domain, internal cross-domain, external Munich, external Frankfurt, internal double-carbapenemase, internal rare-species). Species groups: *Klebsiella pneumoniae* complex, *Escherichia coli, Citrobacter freundii* complex, *Enterobacter cloacae* complex, and others. The final column gives the per-dataset total.

|  |  | *Klebsiella pneumoniae* complex | *Escherichia coli* | *Citrobacter freundii* complex | *Enterobacter cloacae* complex | Others | Total |
| --- | --- | --- | --- | --- | --- | --- | --- |
| Internal in-domain | pos | 67 | 18 | 26 | 12 | 0 | 123 |
|  | neg | 67 | 18 | 26 | 12 | 0 | 123 |
| Internal cross-domain | pos | 16 | 5 | 5 | 4 | 0 | 30 |
|  | neg | 16 | 5 | 5 | 4 | 0 | 30 |
| External Munich | pos | 6 | 3 | 11 | 0 | 0 | 20 |
|  | neg | 7 | 3 | 12 | 0 | 0 | 22 |
| External Frankfurt | pos | 21 | 6 | 5 | 5 | 0 | 37 |
|  | neg | 27 | 14 | 6 | 5 | 0 | 52 |
| Internal double carbapenemases | pos | 22 | 0 | 0 | 1 | 0 | 23 |
|  | neg | 22 | 0 | 0 | 1 | 0 | 23 |
| Internal rare species | pos | 0 | 0 | 0 | 0 | 30 | 30 |
|  | neg | 0 | 0 | 0 | 0 | 30 | 30 |
| Non-IncL OXA-48 isolates | pos | 8 | 22 | 0 | 2 | 0 | 32 |
|  | neg | 8 | 22 | 0 | 2 | 0 | 32 |

**Extended Data Fig. 3.**
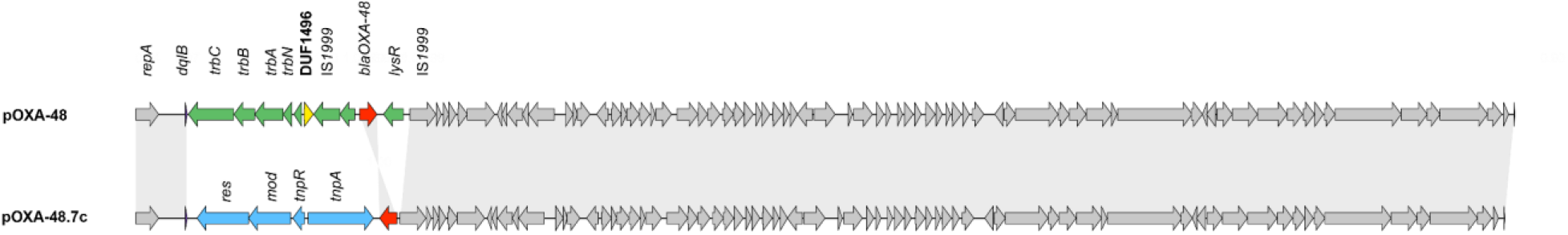
Structural comparison of the reference pOXA-48 plasmid and the naturally occurring variant pOXA-48.7c. Linear gene maps of reference pOXA-48 (NZ_CP124810.1; 63,589 bp; top) and pOXA-48.7c (carried by donor MMC-K-0151, BioSample SAMN25408304; 63,126 bp; bottom). Arrows denote coding sequences in their direction of transcription, with the following colour code: the DUF1496 gene yellow, *bla*OXA-48 red, the conserved plasmid backbone grey, other gene regions unique to pOXA-48 green, and gene regions unique to pOXA-48.7c blue. Shaded bands connect homologous regions. Relative to the reference, pOXA-48.7c has lost the *trb* locus together with the DUF1496 gene.

## Supplementary data

**Supplementary Table 1.** Sensitivity of classification performance to LightGBM hyperparameter settings. Performance of four LightGBM configurations, all using the mixed-protocol, temporally augmented training set-up: the default used throughout (baseline; n_estimators = 200), reduced capacity (n_estimators = 100), increased leaf regularization (min_data_in_leaf = 50), and stochastic regularization (feature_fraction = 0.8, bagging_fraction = 0.8, bagging_freq = 1). AUROC, AUPRC, sensitivity, and specificity are means (s.d.) across the five acquisition protocols and 20 random seeds; TP, FN, FP, and TN are means per-seed counts.

| Metric | AUROC (±s.d.) | AUPRC (±s.d.) | Sensitivity (±s.d.) | Specificity (±s.d.) | TP | FN | FP | TN |
| --- | --- | --- | --- | --- | --- | --- | --- | --- |
| **Baseline (`n_estimators = 200`, defaults)** | 0.963 (0.007) | 0.963 (0.008) | 0.899 (0.022) | 0.892 (0.020) |  | 1.5 | 1.6 | 13.2 |
| **Reduced capacity (`n_estimators = 100`)** | 0.958 (0.008) | 0.958 (0.010) | 0.884 (0.026) | 0.884 (0.019) |  | 1.8 | 1.7 | 13.1 |
| **Increased leaf regularization (`min_data_in_leaf = 50`)** | 0.966 (0.007) | 0.966 (0.008) | 0.909 (0.025) | 0.891 (0.021) |  | 1.4 | 1.6 | 13.2 |
| **Stochastic regularization (`feature_fraction = 0.8`, `bagging_fraction = 0.8`)** | 0.968 (0.007) | 0.968 (0.009) | 0.908 (0.024) | 0.895 (0.017) |  | 1.4 | 1.6 | 13.3 |

**Supplementary Table 2.** Sequencing platforms and library-preparation methods by contributing institution. DNA-extraction kits, short- and long-read library-preparation protocols, and sequencing platforms used for whole-genome sequencing at each contributing institution. Isolates from Hadassah Medical Center, Jerusalem, were sequenced at the University of Zurich. N.a. = not applicable.

| Institution | DNA extraction | Short-read library prep | Short-read platform | Long-read library prep | Long-read platform |
| --- | --- | --- | --- | --- | --- |
| **University Hospital Cologne** | DNeasy UltraClean Microbial Kit (Qiagen) | Illumina DNA Prep | Illumina NextSeq 500 / NovaSeq 6000 | n.a. | n.a. |
| **University of Zurich** | DNeasy Blood & Tissue Kit (Qiagen); Maxwell RSC Blood DNA Kit (Promega) | Qiaseq FX (Qiagen) | NextSeq1000 (PE150) | n.a. | n.a. |
| **Hadassah Medical Center, Jerusalem** | Sequenced at the University of Zurich, see above |  |  |  |  |
| **University** | DNeasy UltraClean | n.a. | n.a. | SQK-LSK109 | GridION; R9.4.1 |
| **Hospital Frankfurt** | Microbial Kit (Qiagen) |  |  | Ligation Sequencing Klt (ONT) with EXP-NBD104/114 Native Expansion kit (R9.4.1) or SQK-BND114.96 Native Barcoding Kit 96 V14 (R10.4.19) | and PromethION P2Solo; R10.4.1 |
| **TU Munich** | DNeasy Blood & Tissue Kit (Qiagen) | n.a. | n.a. | SQK-RBK114 Rapid Barcoding Kit (ONT) | MinION MK1d and PromethION P2Solo; R10.4.1 |
| **Cologne Merheim** | DNeasy UltraClean Microbial Kit (Qiagen) | Illumina DNA Prep | Illumina NextSeq 500 / NovaSeq 6000 | n.a. | n.a. |

**Supplementary Table 3.** Characteristics of the bottom-up proteomics isolates. The 17 clinical isolates analysed by bottom-up proteomics (8 pOXA-48-positive, 9 pOXA-48-negative). For each isolate: isolate ID, species, multilocus sequence type, carbapenemase gene, and IncL plasmid contig size (bp; absent for pOXA-48-negative isolates).

| Isolate ID | Species | Sequence type | Carbapenemase gene | IncL plasmid contig size |
| --- | --- | --- | --- | --- |
| MMC-K-0098 | *Klebsiella pneumoniae* | 15 | OXA-48 | 63585 |
| MMC-K-0051 | *Klebsiella pneumoniae* | 307 | OXA-48 | 63580 |
| MMC-K-0076 | *Escherichia coli* | 131 | OXA-48 | 62785 |
| MMC-K-0255 | *Escherichia coli* | 10 | OXA-48 | 75514 |
| MMC-K-0124 | *Escherichia coli* | 58 | OXA-48 | 62784 |
| MMC-K-0286 | *Enterobacter cloacae* complex | 45 | OXA-48 | 63582 |
| MMC-K-0095 | *Enterobacter cloacae* complex | 114 | OXA-48 | 61574 |
| MMC-K-0194 | *Enterobacter cloacae* complex | 78 | OXA-48 | 62813 |
| MMC-K-0637 | *Klebsiella pneumoniae* | 15 | neg | - |
| MMC-K-0840 | *Klebsiella pneumoniae* | 101 | neg | - |
| MMC-K-0811 | *Klebsiella pneumoniae* | 307 | neg | - |
| MMC-K-0670 | *Escherichia coli* | 131 | neg | - |
| MMC-K-0479 | *Escherichia coli* | 10 | neg | - |
| MMC-K-0808 | *Escherichia coli* | 224 | neg | - |
| MMC-K-0817 | *Enterobacter cloacae* | 815 | neg | - |
| MMC-K-0824 | *Enterobacter cloacae* | 108 | neg | - |
| MMC-K-0819 | *Enterobacter cloacae* | 50 | neg | - |

**Supplementary Table 4.** pOXA-48 peptide precursor matches for the bottom-up proteomics analysis. This supplementary dataset accompanies the journal submission but is not included with this preprint.

**Supplementary Table 5.** Protein-level quantification and predicted masses of the pOXA-48-encoded proteins detected by bottom-up proteomics. This supplementary dataset accompanies the journal submission but is not included with this preprint.

## Notes

### Competing Interest Statement

K.B. is co-founder and scientific advisor of Computomics GmbH, Tuebingen, Germany. A.F.W. has received speaker honoraria from Cepheid. All other authors declare no competing interests.

